# Simulated tempering-enhanced umbrella sampling improves convergence of free energy calculations of drug membrane permeation

**DOI:** 10.1101/2022.11.13.516136

**Authors:** Carla F. Sousa, Robert A. Becker, Claus-Michael Lehr, Olga V. Kalinina, Jochen S. Hub

## Abstract

Molecular dynamics simulations have been widely used to study solute permeation across biological membranes. The potential of mean force (PMF) for solute permeation is typically computed using enhanced sampling techniques such as umbrella sampling (US). For bulky drug-like permeants, however, obtaining converged PMFs remains challenging and often requires long simulation times, resulting in an unacceptable computational cost. Here, we augmented US with simulated tempering, introducing Simulated Tempering-enhanced US (STeUS), to improve the convergence of PMF calculations for the permeation of methanol and three common drug molecules. Simulate tempering helps to enhance sampling by varying the temperature of the system along a pre-defined temperature ladder. To obtain sufficient sampling of the umbrella histograms, which were computed only from the ground temperature, we modified the simulation time fraction spent at the ground temperature between 1/*K* and 50%, where *K* is the number of ST temperature states. We found that STeUS accelerates convergence compared to standard US, and the benefit of STeUS is system-dependent. For bulky molecules, for which standard US poorly converged, the application of ST was highly successful, leading to a more than five-fold accelerated convergence of the PMFs. For the small methanol solute, for which conventional US converges moderately, the application of ST is only beneficial if 50% of the STeUS simulation time is spent at the ground temperature. This study establishes STeUS as an efficient and simple method for PMF calculations, thereby strongly reducing the computational cost of routine high-throughput studies of drug permeability.

## INTRODUCTION

Molecular dynamics (MD) simulations are a powerful computational tool used to obtain an atomistic view of biological processes.^1^ However, studying complex biological systems using MD simulations remains challenging since it requires obtaining exhaustive sampling of the high-dimensional configurational space, which is time-consuming owing to slow conformational dynamics and long autocorrelation times. For instance, simulating systems that include biological membranes at atomic details is challenged by the sticky environment of the head group regions, leading to slow rearrangements of biomolecules or other solutes and, thereby, to slow conformational sampling.^2–5^

Fast and faithful simulation of such systems is, however, very valuable, since solute permeation across biomembranes is highly relevant in the field of drug discovery, as most drugs need to pass multiple cellular membranes to reach their targets. The potential of mean force (PMF, alternatively called the free energy profile) for solute translocation across the membrane can be calculated from MD simulations, providing a spatially resolved view on solute partitioning between water and membrane.^4, 6–8^ For most solutes, membrane permeation is a rare event suggesting that enhanced sampling techniques are needed for computing the PMF (*G*(z)).^9^ Umbrella sampling (US) is one of such tools that has been widely used for computing PMFs. It involves division of the process in windows along a preselected reaction coordinate (RC), while the application of restraining potentials ensures sampling of unfavorable regions of high free energy.^10–14^ These US windows must overlap to sample the complete RC. The PMF is typically obtained by unbiasing the probability distributions from the restrained simulations using the weighted histogram analysis method (WHAM).^15^

For simulations of solute translocation across lipid bilayers, the relative center of mass displacement between solute and membrane along the membrane normal (*z* coordinate) has been often used as RC, thereby sampling the complete permeation event.^13, 16–18^ However, such protocol does not accelerate the convergence of orthogonal degrees of freedom such as the solute orientation, solute–lipid interaction patterns, or the lipid–lipid arrangements. Specifically, simulations of solute insertion in lipid bilayers were found to be affected by the slow reorganization of the ionic interactions between the phospholipid headgroups and the solute.^2, 5, 19^ Slow sampling of such orthogonal degrees of freedom may lead to poor convergence in standard US simulations. Although such problems could in principle be overcome using multi-dimensional US along a set of RCs, finding good RCs is difficult in practice, good RCs may be solute-dependent, and multi-dimensional US is computationally expensive.^20^

Sampling challenges have been addressed by extended-ensemble methods that, in contrast to conventional US, do not require the definition of RCs.^21–26^ One possibility is to use generalized ensemble algorithms in temperature space, such as the simulated tempering (ST) method.^27, 28^ ST aims to achieve broader sampling by periodically changing the temperature during the simulation along a pre-defined temperature ladder. During ST simulations, increased temperatures help the system to overcome enthalpic barriers and, thereby, to enhance the conformational sampling. Importantly, unlike simulated annealing that drives the system out of equilibrium, ST maintains equilibrium and produces well-defined thermodynamic ensembles. Hybrid methods that combine reaction coordinate-based with extended ensemble-based methods have also been proposed.^29–33^

I In this study, we introduce a new protocol for combining ST with US (ST-enhanced US, or STeUS) and analyze its benefit for the calculation of PMFs for solute permeation, by comparing the convergence of the PMFs between standard US and STeUS. To cover solutes of different molecular weights, area/volume and polarity, we simulated the permeation of methanol, ibuprofen, 1-propranolol and atenolol (Table 1). Except for methanol, these molecules are widely used drugs with common physicochemical properties, suggesting that our results are representative for future simulations in drug development. We show that STeUS is a promising approach for improving the efficiency and accuracy of free energy calculations of membrane permeation.

**Table 1.**
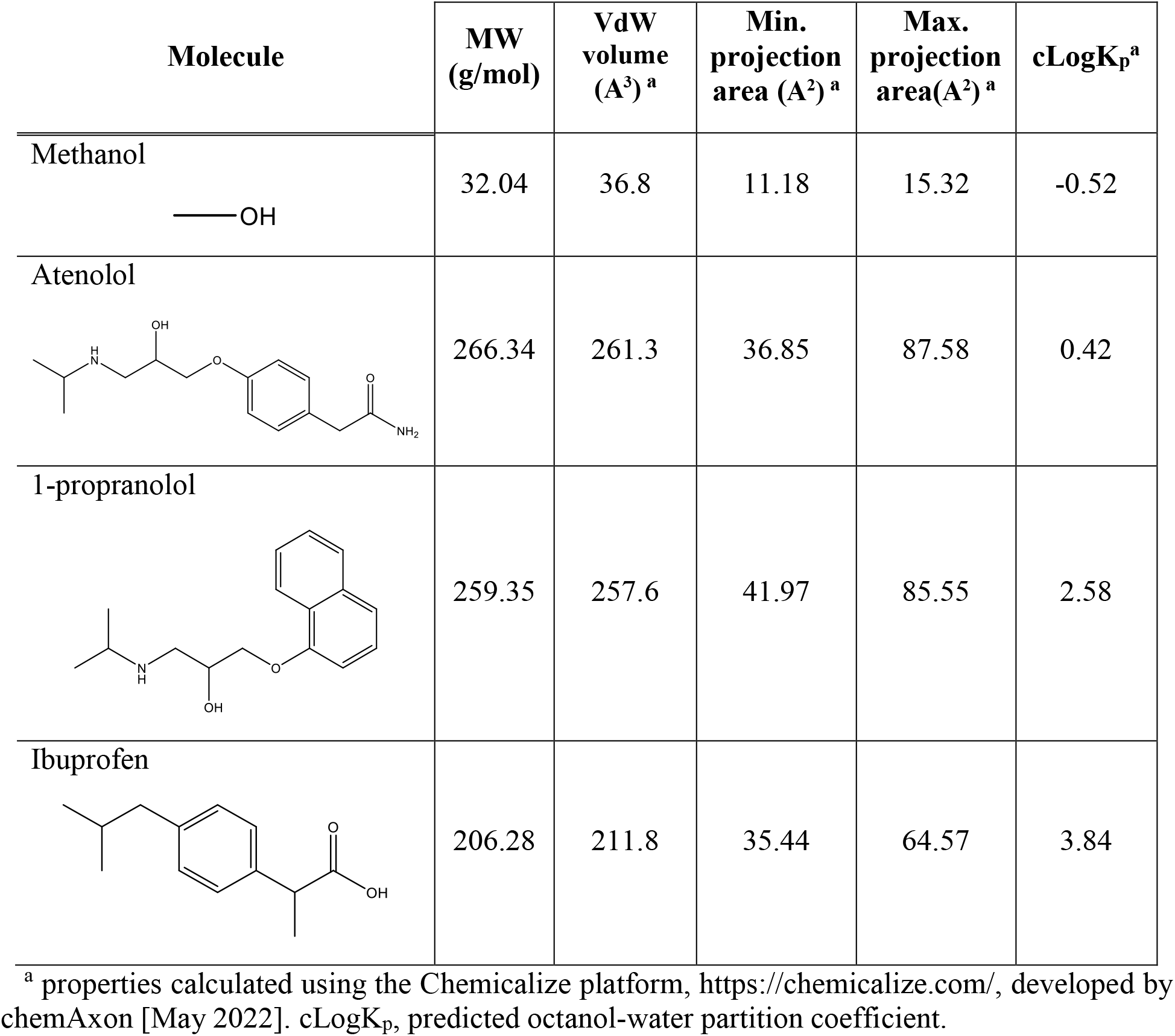
Physicochemical properties of the molecules used in this study.

| Molecule | MW<br>(g/mol) | VdW<br>volume<br>(Å<sup>3</sup>)<sup>a</sup> | Min.<br>projection<br>area (Å<sup>2</sup>)<sup>a</sup> | Max.<br>projection<br>area(Å<sup>2</sup>)<sup>a</sup> | cLogK<sub>p</sub><sup>a</sup> |
| --- | --- | --- | --- | --- | --- |
| Methanol | 32.04 | 36.8 | 11.18 | 15.32 | -0.52 |
| Atenolol | 266.34 | 261.3 | 36.85 | 87.58 | 0.42 |
| 1-propranolol | 259.35 | 257.6 | 41.97 | 85.55 | 2.58 |
| Ibuprofen | 206.28 | 211.8 | 35.44 | 64.57 | 3.84 |

<sup>a</sup> properties calculated using the Chemicalize platform, https://chemicalize.com/, developed by chemAxon [May 2022]. cLogK<sub>p</sub>, predicted octanol-water partition coefficient.

In ST, weights are applied to control the occupancies of the *K* temperature states (*T*_0_ … *T*_*K*-1_), while these weights are often optimized to obtain uniform occupancies among the temperature states. For US, however, uniform occupancies may not be optimal, because only a fraction of 1/*K* of the simulation time is available for collecting the umbrella histograms at the physiologically relevant ground temperature *T*_0_. Therefore, we devised a modified STeUS protocol that allows an increased, user-defined, occupancy *P*_0_ of the ground temperature *T*_0_, while all higher temperature states remain uniformly populated. As shown below, the benefit of an increased occupancy of the ground temperature is dependent on the solute’s physicochemical characteristics, especially its size. We found that for small molecules such as methanol, the simulation time spent at *T*_0_ may limit the convergence of the umbrella histograms, suggesting the use of higher values of *P*_0_. For bulkier, drug-like solutes, in contrast, longer simulation times at higher temperatures were most critical for convergence, suggesting that only a moderately increased *P*_0_ is most beneficial for obtaining rapid sampling.

## THEORY

### Simulated Tempering

In ST, the temperature is a dynamic variable, and a random walk is performed on the temperature ladder of (*T*_0_ … . *T*_*K*-1_).^27, 34^ The appropriate choice of the temperature ladder is important: *T*_0_ typically corresponds to the physiologically relevant temperature while *T*_*K*-1_ should be high enough to aid the transition over relevant free energy barriers.

Let *x* denote the configuration and *n* = 0, …, *K* – 1 the temperature state, then the probability distribution *P*(*x, n*) is:

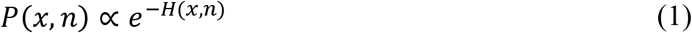

where the generalized Hamiltonian *H*(*x,n*) is defined by:

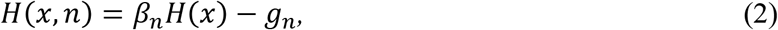

with *β_n_* = 1/*k_B_T_n_* being the inverse temperature and *g_n_* the weight for the *n*th temperature.

Hence, the partition function is

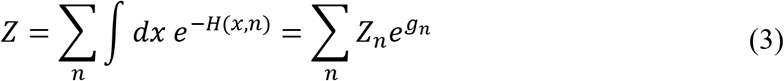

where *Z_n_* is the partition function for state *n* with temperature *T_n_*. The Hamiltonian implies a generalized ensemble with canonical ensembles for each temperature, while relative weights of the temperatures are modified with the parameters *g_n_*.

Typical ST simulations involve common MD simulations at a constant temperature *T_n_* with updates to neighboring temperatures *T*_*n*+1_ or *T*_*n*−1_. Transitions between *T_n_* and *T*_*n*±1_ are performed according to the Metropolis criterion with a probability^35^

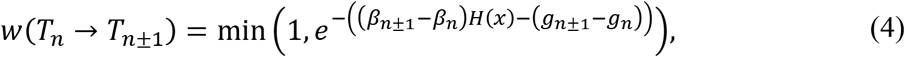

thereby maintaining detailed balance and well-defined canonical ensembles. During a burn-in phase at the beginning of the simulation, the weights *g_n_* are typically optimized to obtain uniform occupancies of all states.

The burn-in phase is greatly simplified by choosing good initial weights, as suggested by Park *et al*.^36^ Accordingly, requesting that optimal weights yield equal acceptance for transition *T_n_* → *T*_*n*+1_ and *T*_*n*+1_ →*T_n_*, we have Δ*H*_*n*→*n*+1_ = Δ*H*_*n*+1→*n*_ with

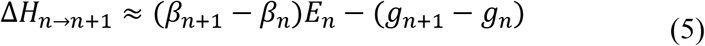

and

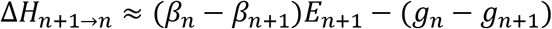

Hence, the relative weights can be calculated from:

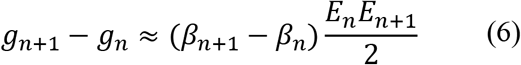

where *E_n_* is the average potential energy of the system at *T_n_*, which can be taken from a short simulation.

### Increased occupancy of the ground temperature

In this study, the weights were updated with the Wang-Landau algorithm.^37^ This approach involves updating the weights (*g_n_*) by an incrementor factor (*δ*) with the goal to obtain uniform histogram over the temperature states. During a burn-in phase, the weights are optimized in rounds, where each round involves a random walk over the temperature states using a given set of initial *g_n_*. In every state transition, *g_n_* is updated by *g_n_new__* = *g_n_old__* × *δ*, until a flat histogram over temperature is achieved, indicating the convergence of the *g_n_* with an accuracy proportional to ln(*δ*). In the next round, *δ* is reduced and a new random walk over the states is started. As the simulation proceeds, these steps are repeated, while *δ* decreases monotonically to 1, leading to an increasingly accurate convergence of the *g_n_^3^*

Whereas the traditional ST approach results in uniform occupancy over temperatures (*T*_0_ … *T*_*K*-1_), only the samples generated at the groumd temperature *(T*_0_) are typically physiologically relevant.^38^ This is true for the specific case of PMF calculations for drug membrane permeation: while sampling at higher energies is essential to overcome free energy barriers, only a small fraction of 1/*K* of samples at the ground temperature are correctly distributed to allow the construction of the PMF for the physiologically relevant temperature. To accelerate the convergence of the umbrella histograms at *T*_0_, we devised a modified ST approach that spends an increased, user-defined, occupancy *P*_0_ (instead of 1/*K*) at the ground temperature. This was implemented with a reduced Wang-Landau incrementor factor *δ*_*P*_0__ applied for the ground temperature. Specifically, let *P_h_* denote the occupancy of the higher states, then we have *P*_0_ + (*K* − 1)*P_h_* = 1. Hence, *δ*_*P*_0__ was reduced by a factor of *P_h_/P_0_* for the ground temperature, while the incrementor for other states was chosen following the usual Wang-Landau method. A modified source code file expanded.cpp for GROMACS 2021 that implements the methods is available as supporting information.

## METHODS

The membrane system, comprising 32 palmitoyl-oleoyl-phosphatidylcholine (POPC) phospholipids and 50 water molecules per lipid in a cubic simulation box, was taken from Nitschke *et al*.^20^. Lipids were described using the modified Berger force field, as defined by Nitschke *et al*.^20, 39, 40^ and water by the SPC model^41^. Methanol was modelled using the Gromos43A1 force field and the distance between the carbon and the hydroxyl hydrogen was constrained to allow the use of a 5 fs time step. Ibuprofen, atenolol and 1-propranolol were modelled using the Gromos54A7 force field, with parameters from the Automated Topology Builder (ATB) website^42^. Since this study focuses purely on sampling, we did not further validate the force fields against experimental data.

MD simulations were performed and analyzed using GROMACS 2021^43, 44^ and visualized using VMD^45^. The center of mass (COM) motion of the bilayer relative to the solvent was removed every 10 integration steps. COM groups were: (i) each lipid monolayer, and (ii) solvent and ligands. The Particle-mesh Ewald (PME) method^46, 47^ was used for long-range electrostatic interactions, with a real space cut-off of 1.2 nm, and a Fourier spacing of 0.14 nm and a 1.2 nm cut-off was used for the Van der Waals interactions. Long-range dispersion corrections to pressure and energy were applied^48^. The v-rescale thermostat^49^, with 0.5 ps coupling constant, was used for temperature coupling and the Berendsen barostat^50^, with 10 ps coupling constant, for pressure coupling. The LINCS algorithm was used to constraint all bonds of lipids and solutes^51^. Water molecules were constrained with SETTLE^52^.

Drugs were inserted into the system, as previously described^13^: two drugs molecules were inserted into the system (one into water, *z* = −3.5 nm, and the other at the bilayer center, *z* = −0.2 nm) by gradually switching on their Van der Waals and Coulombic interactions with the system, using ten λ steps (0.6 ns each). The COM position of the solute relative to the bilayer along the bilayer normal (*z*) was used as the reaction coordinate for US simulations.^10^ Here, only lipid atoms within a cylinder of radius 1.5 nm around the solute were used to compute the bilayer COM, as implemented in the pull geometry “cylinder” of the GROMACS code. Initial structures for each of the 37 US windows were generated using sequential 5 ns simulation, where the solutes were pulled over a distance of Δ*z* = 0.1 nm in each simulation. Each US window was simulated for 100 ns, using a harmonic force constant of 1000 kJ·mol^-1^·nm^-2^, and the productions were repeated five times. The first 10 ns were removed for equilibration and PMFs were constructed using WHAM.^15, 53^ Five replications were performed for each system tested in this study. Errors of the profiles for each individual replica were estimated using the Bayesian bootstrap analysis of complete histograms, using a total of 200 bootstraps.^53^ Errors of the profiles for the average of the 5 replicas, represent the standard deviation between these data.

For the simulated-tempering protocol the md-vv integrator was used, a velocity Verlet algorithm for integrating Newton’s equations for motion. Nine temperatures were used (*T*_0_ = 300 K … *T*_8_ = 348 K) with a Δ*T* = 6 K. Initial weights were determined as described by Park *et al.^36^.* Sequential short simulations (10 ps) were performed for each temperature (*T*_0_ … *T*_8_), and the weights were determined from the average potential energy, at each temperature.

Two values of target occupancy of the ground temperature (*P*_0_) were tested: according to the standard ST protocol, *P*_0_ = 1/9, as well as, according to the modified ST protocol, *P*_0_ = 0.2 or *P*_0_ = 0.5 (Figure 1).

**Figure 1.**
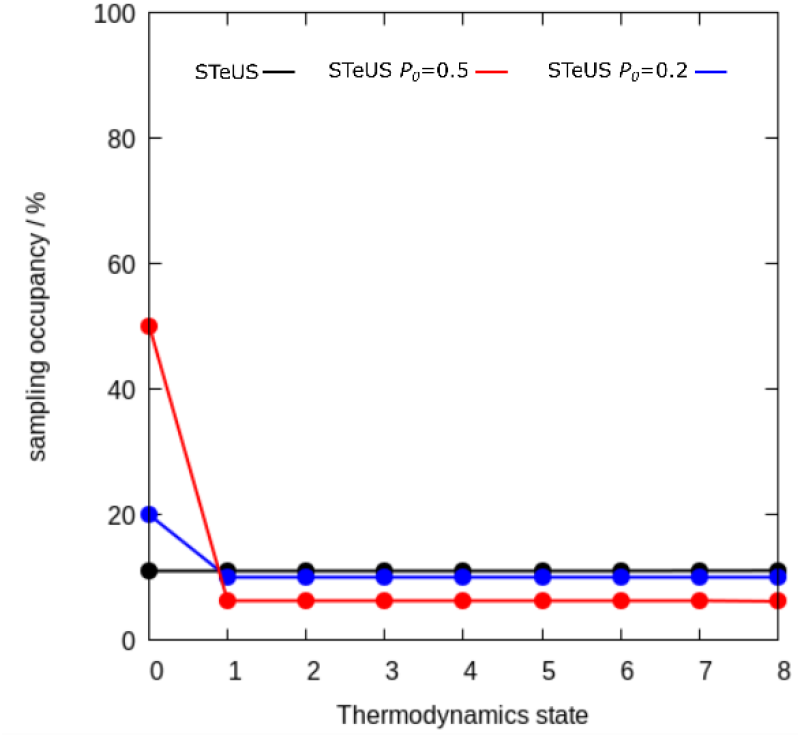
Occupancy of temperature states (*T*_0_ = 300 K, …, *T*_8_ = 300 K; Δ*T* = 6 K) in simulations with ibuprofen for STeUS with uniform occupancies (black) or with increased occupancies of *P*_0_ = 0.2 and *P*_0_ = 0.5 for the ground temperature (blue and red, respectively).

## RESULTS

### Standard umbrella sampling leads to poor convergence of PMFs for drug-like permeants

PMFs of solute translocation across the POPC bilayer were obtained with traditional US, comprising 100 ns simulation time per window, while omitting 10 ns for equilibration, being each window simulated 5 times (Figure 2 and S1). PMFs were computed for solute translocation from bulk water into the membrane center (inward direction) and from the membrane center to bulk water (outward direction). Because the bilayer is symmetric, converged PMFs of the two molecules should likewise be symmetric. Hence, in this work we used the absolute difference between the PMFs for inward and outward directions as a measure for the convergence of the PMFs (Figure S2). Considerable free energy offsets between the PMF for the inward and outward directions are evident for all bulkier drug-like permeants - atenolol, 1-propranolol and ibuprofen, - indicating major hysteresis problems and poorly converged PMFs. This conclusion is also supported by a large difference between the PMFs of the 5 independent replicas (Figure S1 and Figure 2). At the same time, PMFs for the smaller permeant – methanol - show a much better convergence and less hysteresis (Figure 2).

**Figure 2.**
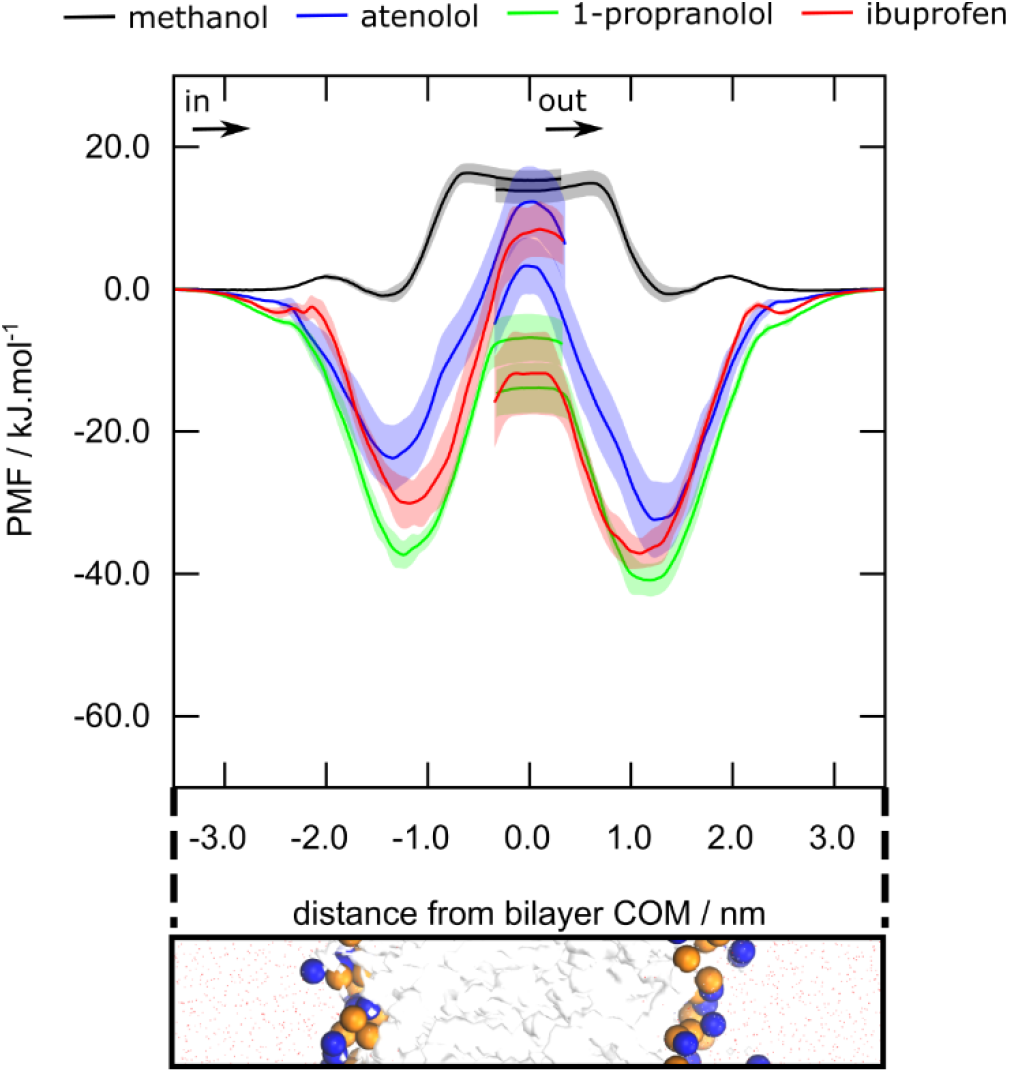
PMFs for methanol (black), atenolol (blue), 1-propranolol (green), and ibuprofen (red) as a function of distance to the bilayer COM at *z* = 0.0 nm, based on 5 independent replicas for each solute and 100 ns of simulated time, using standard US. Shaded areas represent the standard deviation between the replicas. For reference, the lower panel shows the bilayer with water molecules depicted as red dots, membrane as a white surface, and phosphate and choline groups as orange and blue spheres, respectively.

To test whether longer simulation times reduce the hysteresis error, we followed the PMFs using increasing simulation times of 20 ns to 100 ns in steps of 10 ns (Figure 3). However, in this simulation time range, longer simulations reduced the hysteresis error only partly, suggesting that the hysteresis was caused by autocorrelations on longer time scales and that by far longer simulations would be needed to obtain converged PMFs with conventional US.

**Figure 3.**
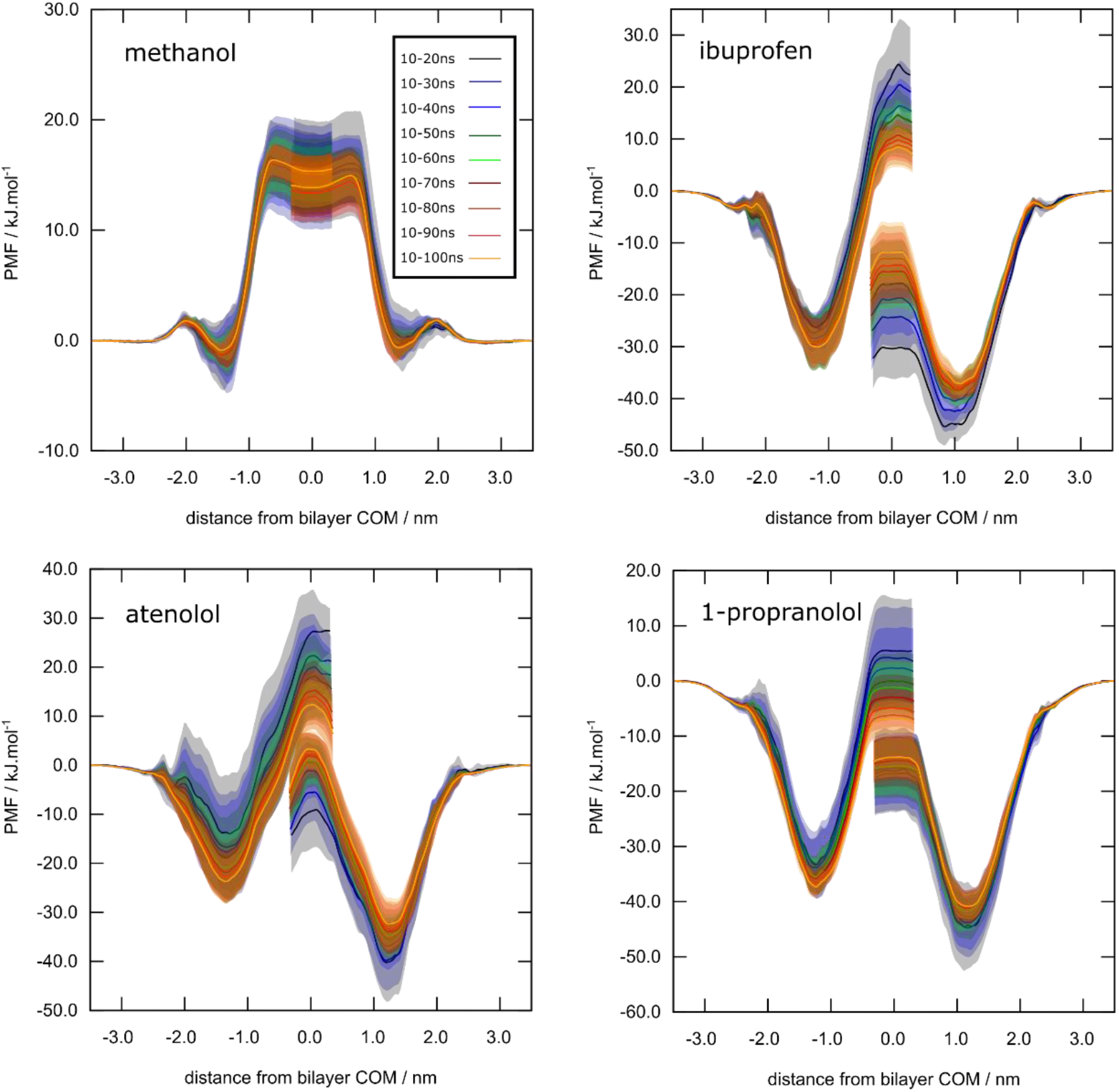
PMF for each solute, methanol, ibuprofen, atenolol and 1-propranolol, with increasing simulation times of 20 ns to 100 ns in steps of 10 ns, using conventional US. Shaded areas represent the standard deviation between the 5 independent replicas.

Statistical errors for each replica (Figure S1, shaded areas) were estimated by bootstrapping complete histograms. Such approach typically yields a conservative error estimate because it assumes that only different histograms are independent, but it does not assume a specific autocorrelation time within individual US windows. As evident from Figure S1, however, the statistical error dramatically underestimates the true error owing to hysteresis. This demonstrates that neighboring histograms are highly correlated, and reinforces that the 100 ns of conventional US simulations were, by far, insufficient to faithfully reproduce PMFs.

### Simulated tempering-enhanced umbrella sampling (STeUS) accelerates the convergence of PMFs

Simulated tempering was integrated in the previous US protocol employing 9 temperature states from 300 K to 348 K in steps of ΔT = 6 K, yielding a new simulated tempering-enhanced umbrella sampling (STeUS) protocol. Using it, the PMF at the ground temperature of 300 K was calculated (Figure 4 and S3), equivalent to the temperature adopted during the conventional US simulations (Figure 2 and S1).

**Figure 4.**
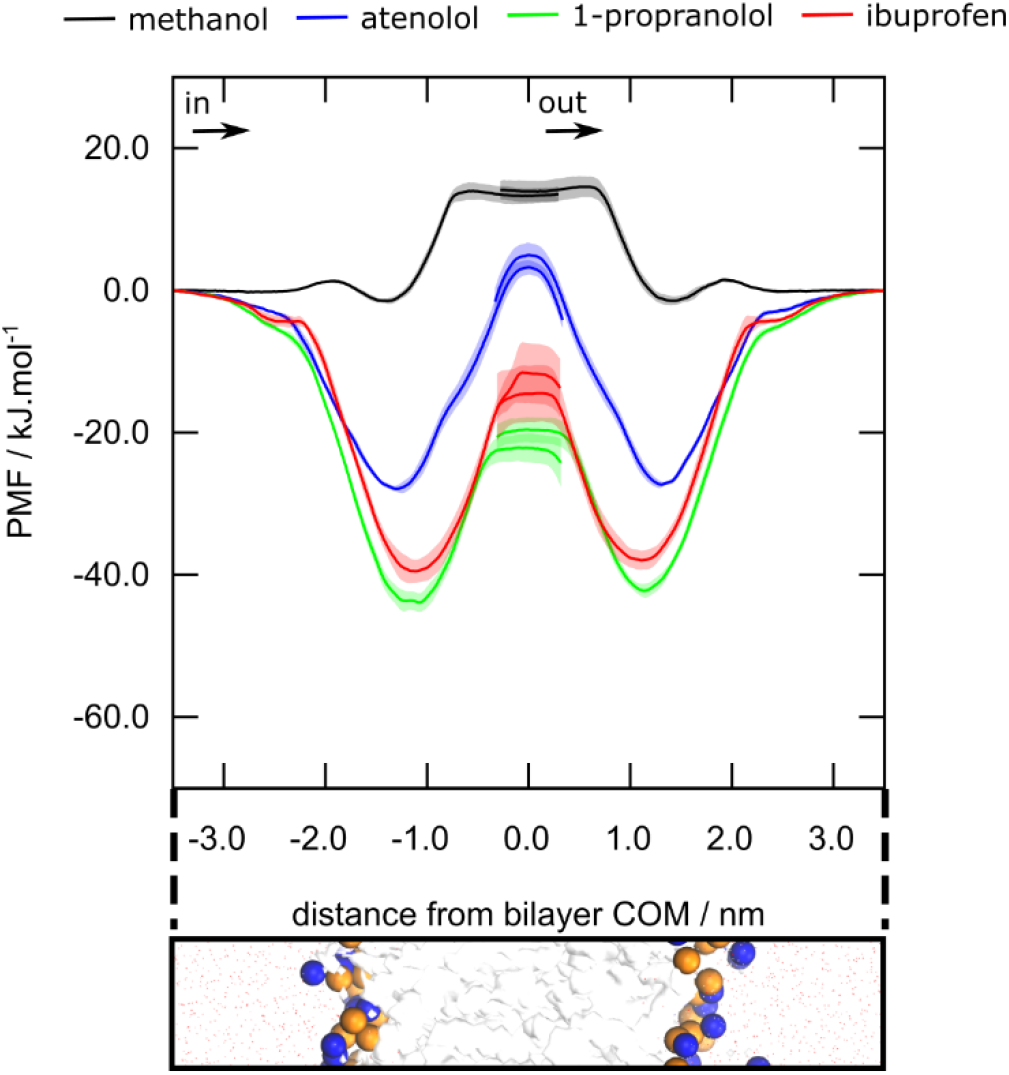
PMFs for methanol (black), atenolol (blue), 1-propranolol (green), and ibuprofen (red) as a function of distance to the bilayer COM at *z* = 0.0 nm, based on five independent replicas for each solute and 100 ns of simulated time using the STeUS method. Shaded areas represent the standard deviation among the replicas. For reference, the lower panel shows the bilayer with water molecules depicted as red dots, membrane as a white surface, and phosphate and choline groups as orange and blue spheres, respectively.

The application of STeUS for the bulkier drug-like solutes atenolol, 1-propranolol and ibuprofen led to a marked decrease in the offsets between the PMFs for the inward and outward directions (Figure 4 and Figure S3), demonstrating a greatly decreased hysteresis error for the bulky solutes (Figure 5). In addition, the PMFs were more symmetric regarding the shapes and positions of the PMF maxima and minima, and the variability among the five independent replicas was considerably decreased as compared to the standard US. These observations indicate greatly improved convergence of the PMFs within the time frame of the study with STeUS (Figure S3). Moreover, PMFs computed with increasing simulation times, in steps of 10 ns, from 20 to 100 ns indicate that, unlike US, the STeUS method yields reasonable convergence within short simulation times of 20 ns to 50 ns depending on the solute (Figure S4).

**Figure 5.**
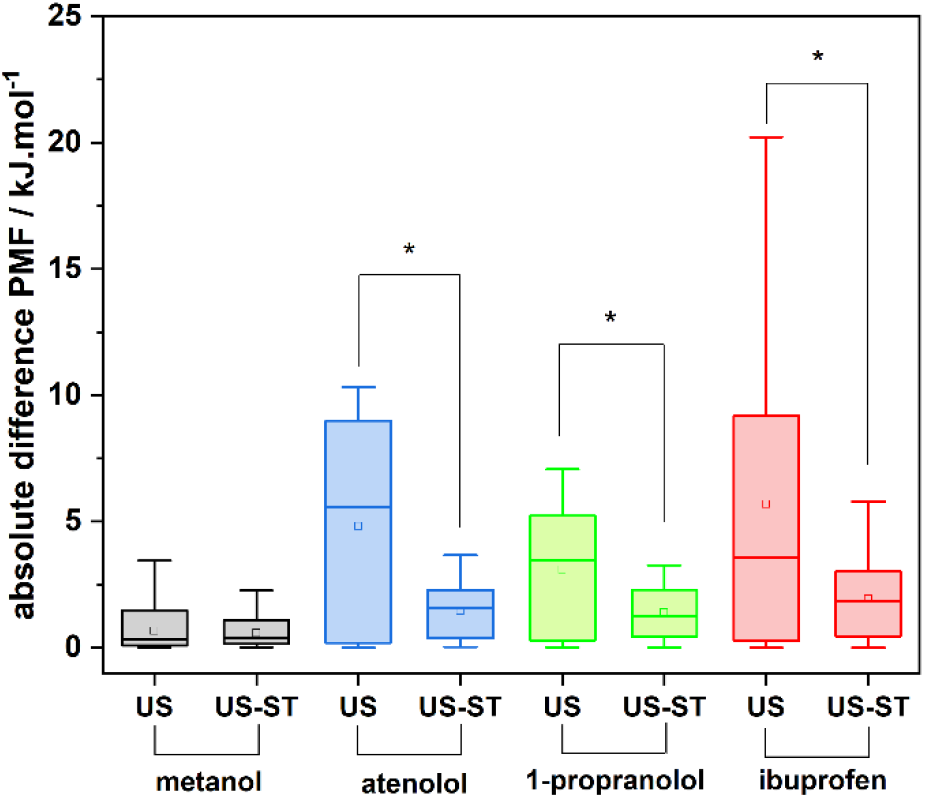
Boxplot representation of the absolute difference between the PMFs for the molecule in the inward and outward directions relative to the membrane, after 100 ns of simulation. Data from the US (Figure 2) and the STeUS (Figure 4) approaches were compared using the t-test (stars represent p < 0.0001).

The impact of the improved convergence of the PMFs on the determination of macroscopic thermodynamic properties, which can be compared with experimental quantities, was assessed by calculating the partition coefficient (LogKp) of the solutes to the model membrane after 30 and 100 ns of simulated time (Table S1). Noteworthy, application of the STeUS approach significantly reduced the error of LogKp, being the error of the determination already minimal after 30 ns of simulation and lower than the error of US after 100 ns. Further indication of faster convergence is the greatly reduced difference between the LogKp for the molecules in the inward and outward directions obtained with STeUS (Table S2).

### Increasing the occupancy of the ground temperature further accelerates the convergence of PMFs in a solute-dependent manner

The results reported above indicate that the enhanced convergence obtained by integrating ST in US is solute dependent. While the decrease in hysteresis error for the bulkier solutes was greater, for the small methanol STeUS only marginally reduced hysteresis (t-test p = 0.168). Additional insights in the convergence is given by analyzing the average absolute difference between the PMFs for the molecules in the inward and outward directions as a function of simulation time, which gives a quantitative analysis of the hysteresis error (Figure 6). For all the bulkier permeants, atenolol, ibuprofen and 1-propranolol, there is a substantial advantage in using the STeUS approach. The hysteresis in the 20 ns simulation is lower than the one of the conventional US method after 100 ns of simulation, suggesting a more than 5 times increased convergence for the STeUS protocol (Figure 6B, C and D). For methanol, however, the benefit of using STeUS is only marginal and starts to be evident only after 60 ns of simulation (Figure 6A). In fact, for this molecule, the hysteresis error is very small for both the US or STeUS approaches, with the average absolute difference for the PMFs below 2 nm after 30 ns of simulation. This can be explained by the small size of methanol, for which the convergence is much faster as compared to bulkier permeants.

**Figure 6.**
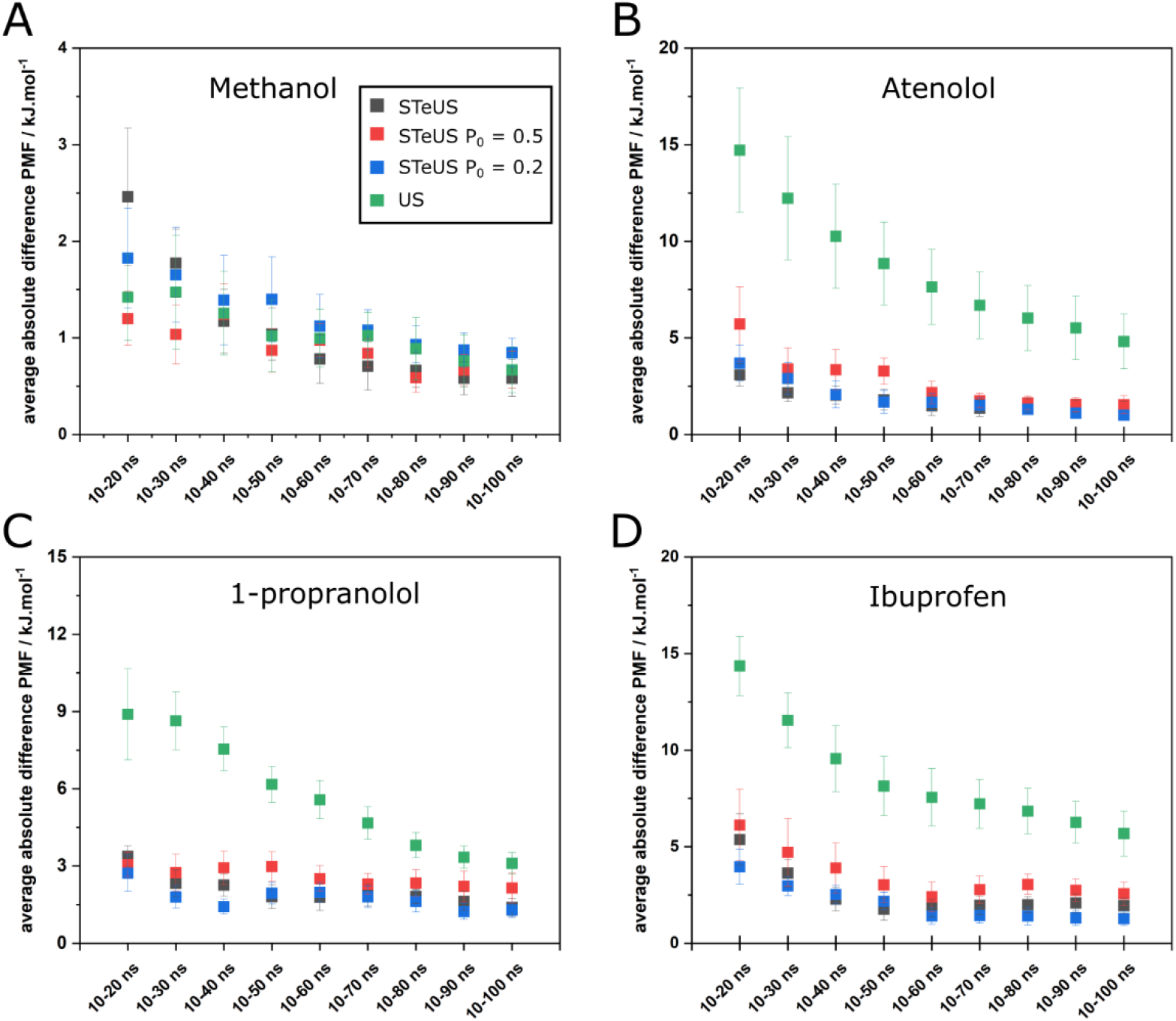
Average absolute difference between the PMFs for the molecule A) Methanol; B) Atenolol; C) 1-propranolol; D) Ibuprofen, in the inward and outward directions, with increasing simulation times. The error in each point represents the standard error between the 5 independent replicas. Data points for each approach used (US, STeUS and modified STeUS with P_0_ = 0.2 and P_0_ = 0.5) are depicted with different colors.

By applying the STeUS approach, in which the system is sampled equally between the different temperatures used (*T*_1_ = 300 K … *T*_9_ = 348 K, Δ*T* = 6 K), the time spent at the ground temperature (300 K) is only approximately 11% of the total simulation time (Figure 1). Hence, since only the frames obtained from the simulation at 300 K were used to collect the umbrella histograms and, thereby, to compute the PMFs, errors also arose from insufficient simulation time at the ground temperature (Figure S4). These errors seem to affect specially the performance of STeUS, compared to US, for the smallest permeant methanol. In an attempt to circumvent this problem, we implemented a modified ST protocol into GROMACS that allows defining the fraction *P*_0_ of simulation time spent at the ground temperature (see Theory). As a control, applying the modified STeUS approach with *P*_0_ = 11% yields similar results as standard STeUS because the occupancy of the ground state is similar (Figures S5 and S6).

Using the modified STeUS method, we computed PMFs with *P*_0_ = 20% or *P*_0_ = 50% (Figure 6). For methanol, *P*_0_ = 50% leads to a slightly reduced hysteresis for short simulation times (Figure 6A, 10-20 ns). Using *P*_0_ = 20% does not yield any improvement for methanol compared to the standard STeUS. On the other hand, for the bulkier permeants, atenolol, 1-propranolol and ibuprofen, using 50% occupancy at the ground temperature leads to a worse performance than the standard STeUS approach (Figure 6B, C and D). Conversely, doubling the sampling at the ground temperature (*P*_0_ = 0.2) leads to a small improvement in convergence, indicating that this might be the optimal occupancy for bulkier permeants (Figure 6B, C and D).

## DISCUSSION

We analyzed the convergence of PMF calculations for membrane permeation based either on conventional umbrella sampling (US) simulations or on US augmented with simulated tempering (STeUS). Four solutes with different physicochemical characteristics were tested: methanol, atenolol, 1-propranolol and ibuprofen (Table 1). Our analysis suggests that PMF calculations for bulky drug-like compounds converge poorly when using conventional US, whereas combining US with ST (STeUS) greatly accelerates the convergence of the PMFs by at least five-fold (Figures 5 and 6). The improved convergence with STeUS was furthermore evident from the greatly reduced statistical error of the membrane partition coefficient for these solutes. In contrast, for small compounds such as methanol, conventional US performs well, suggesting that STeUS does not provide a benefit.

This study focused on sampling with the aim to obtain converged PMFs for a given force field and we did not aim to refine the force fields against experimental data. For the interest of the reader we note, however, that the computed water/membrane partition coefficients of the three bulkier molecules deviate significantly from experimental values (Table S1).

While potentially providing the benefit of getting over the energetic barriers while crossing the membrane, the conventional implementation of ST may lead to slower convergence of the umbrella histograms because only a fraction of 1/*K* of simulation time is spent at the physiologically relevant ground temperature, where *K* is the number of temperature states (Figure 1). To mitigate this limitation, we devised a modified ST implementation that allows the definition of the simulation time fraction *P*_0_ spent at the ground temperature *T*_0_. Specifying *P*_0_ enables finding a good balance between (i) simulating at *T*_0_ to collect samples for the umbrella histograms and (ii) simulating at higher temperatures to overcome sampling barriers more rapidly. We found that an optimal choice of *P*_0_ depends on the solute characteristics.

For the bulkier solutes, extensive sampling at higher temperatures was more critical and spending *P*_0_ = 11% of the simulation time at *T*_0_ was sufficient for obtaining at least five-fold accelerated convergence. Increasing the occupancy of the ground temperature to *P*_0_ =20% further improved the convergence by a minor margin. For the small solute, methanol, for which the PMFs converged even with conventional US, STeUS achieved faster convergence only when spending *P*_0_ = 50% at the ground temperature. Since (i) pharmaceutically relevant solutes are bulky, while (ii) PMF calculations with small solutes converge reasonably rapidly irrespective of the computational details, we suggest a value of *P*_0_ = 20% as a useful starting point for future simulations.

For the small systems tested in this study, the use of nine temperature states was sufficient to obtain a significant improvement in convergence using STeUS. For bigger systems, however, a broader range of temperature states will be necessary, decreasing the percentage of time spent at the physiologically relevant ground temperature. In this case, the modified ST implementation will bring a greater improvement to the sampling of the umbrella histograms.

The use of ST may lead to a moderate increase of computational cost as compared to MD because of the requirement of specific integrators. With GROMACS 2022, ST requires the use of the velocity Verlet integrator instead of the leap-frog algorithms leading to a loss of computational efficiency by ~20% for the systems considered in this study. For the PMF calculations of bulkier compounds, this loss is by far outweighed by the convergence improvement, which is at least fivefold.

ST was previously used to enhance conventional MD simulations, for example, in the context protein folding.^54, 55^ To the best of our knowledge, this work for the first time combines ST with US and employs ST for studying solute permeation. Alternative approaches, such as replica exchange methods (*e.g*. Hamiltonian replica exchange) were previously demonstrated to improve convergence for membrane-solute permeation studies.^56^ A disadvantage of replica exchange methods is, however, that simulation replicas must be conducted in parallel, whereas STeUS allows for a trivial parallelization with fully independent umbrella windows. Furthermore, Rauscher *et al*.^25^ suggested that ST-based methods are more efficient than replica exchange-based methods for sampling complex conformational landscapes. They also argued that, given that accurate weights are obtained, ST was most efficient among the generalized-ensemble methods tested in their study.

## CONCLUSION

We showed that the application of the simulated tempering-enhanced umbrella sampling (STeUS) accelerates convergence of PMFs (or free energy profiles) for membrane permeation of drug-like molecules as compared to the conventional umbrella sampling. The degree of improvement provided by STeUS was solute dependent. STeUS was particularly successful for bulky drug-like molecules, providing an at least five-fold increase of sampling efficiency. For smaller permeants, the application of simulated tempering was only beneficial if the system spends 50% of the simulation time at the physiologically relevant ground temperature. The option to define the proportion of time spent at the ground temperature was implemented in the modified ST protocol developed in this study. We expect STeUS to help establishing all-atom MD simulations and PMF calculations as a high-throughput method for predicting membrane permeability of drug candidates.

## Supporting information

STeUS modified code

## Author Contributions

The manuscript was written through contributions of all authors. All authors have given approval to the final version of the manuscript.

## ACKNOWLEDGMENTS

C.F.S. gratefully acknowledges financial support by the “Singh-Chhatwal-Postdoctoral Fellowship Program”, of the “Friends of HZI” association. O.V.K. acknowledges the funding by the Klaus Faber Foundation. R.B. and J.S.H were supported by the Deutsche Forschungsgemeinschaft (DFG, German Research Foundation, grant no SFB 860/A16).

## ABBREVIATIONS

US: umbrella sampling
ST: simulated tempering
STeUS: simulated tempering enhanced umbrella sampling
MD: molecular dynamics
PMF: potential of mean force
T: temperature
WHAM: weighted histogram analysis method
POPC: palmitoyl-oleoyl-phosphatidylcholine
COM: center of mass.

## Notes

### Competing Interest Statement

The authors have declared no competing interest.

