## supporting Information for "Simulated tempering-enhanced umbrella sampling improves convergence of free energy calculations of drug membrane permeation"

#### Standard umbrella sampling leads to poor convergence of PMFs for drug-like permeants

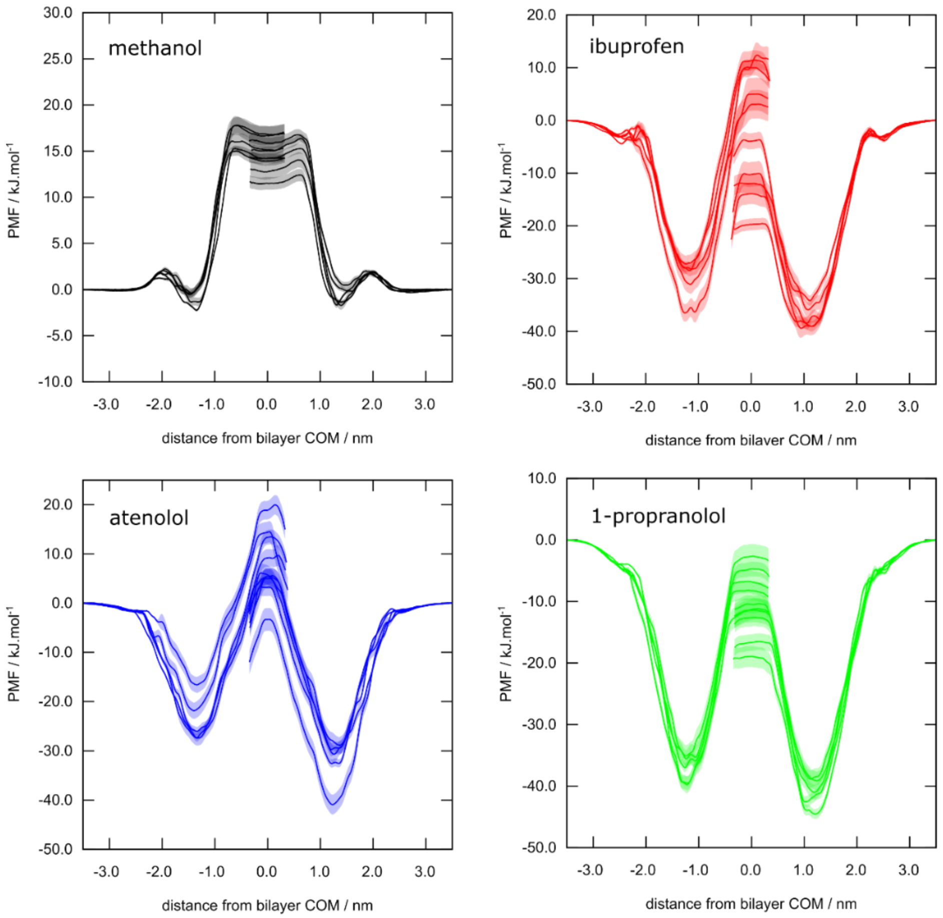


**Figure S1.** PMF as a function of solute distance from the bilayer COM, z = 0.0 nm, using standard US. The multiple solid lines represent the 5 replicas performed for each permeant. Dashed areas represent the statistical errors for each replica, estimated by bootstrapping.

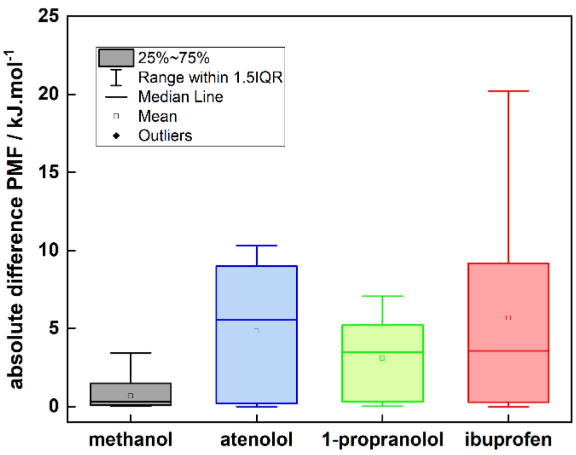


**Figure S2.** Boxplot representation of the absolute difference between the PMF for the molecule in the inward and outward directions relative to the membrane, revealing large undesired hysteresis effects.

### Simulated tempering-enhanced umbrella sampling accelerates the convergence of PMFs

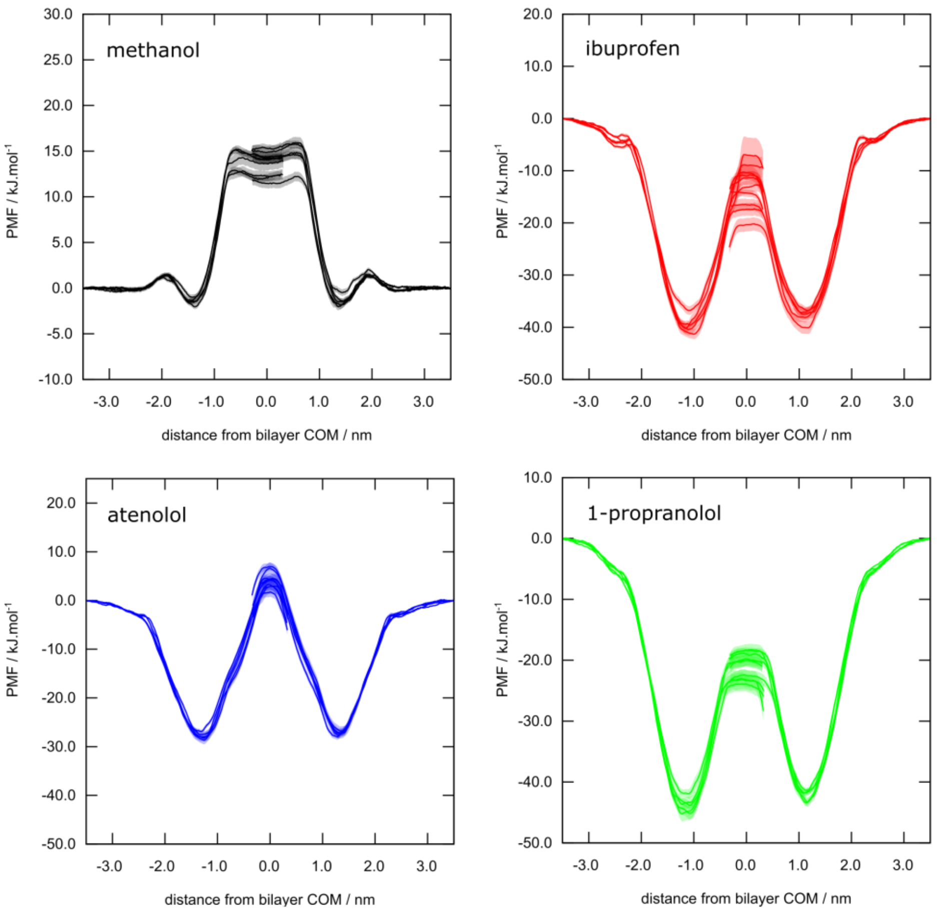


**Figure S3.** PMF as a function of solute distance from the bilayer COM, z = 0.0 nm, using STeUS. The multiple solid lines represent the 5 replicas performed for each permeant. Dashed areas represent the statistical errors for each replica, estimated by bootstrapping.

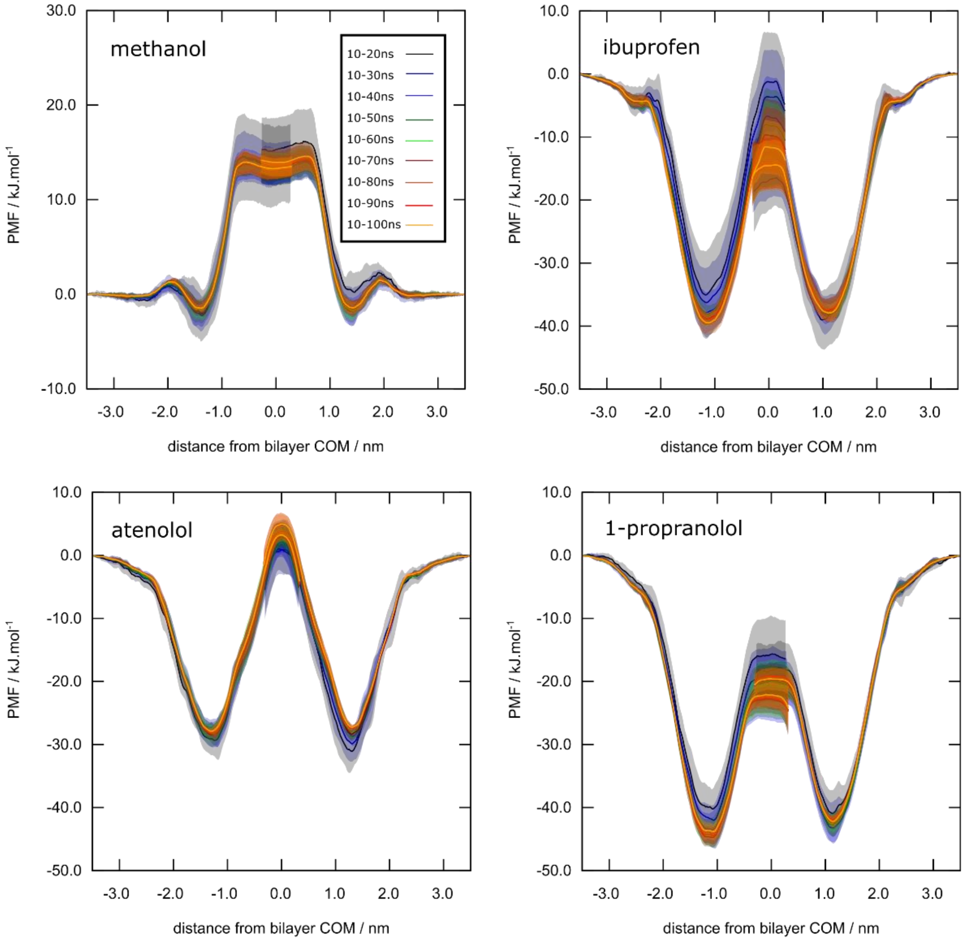


**Figure S4.** PMF for each solute, methanol, ibuprofen, atenolol and 1-propranolol, with increasing simulation times of 20 ns to 100 ns in steps of 10 ns, using the STeUS approach.

### Application of STeUS improves partition coefficient determination

The partition of the solute to the membrane phase ($Kp_{w\rightarrow m}$) can be determined from its binding free energy to the membrane ($\Delta G^0_{bind}$):[1-3]

$$\Delta G^0_{bind} = -k_B T ln\left(\frac{1}{2z_b}\int_{-z_b}^{z_b} e^{-\frac{1}{k_B T}G(z)}\, dz\right) \quad (1)$$

$$K_p = e^{-\frac{1}{k_B T}\Delta G^0_{bind}} \quad (2)$$

where $k_B$ is the Boltzmann constant and $z_b$is the absolute distance of the solute from the bilayer center where it reaches bulk water, and where the PMF is defined to zero.

**Table S1.** The partition constant, expressed as $logK_p \pm SD$, for each molecules is this study. Comparison between experimental and computational values obtained with US and STeUS, after 30 and 100 ns of simulated time.

| Molecule | $LogK_p$ exp. | $LogK_p$ comp. 10 – 30 ns | | 10 – 100 ns | |
|---|---|---|---|---|---|
| | | US | STeUS | US | STeUS |
| **Methanol** | -0.69[a] | -0.198±0.008 | -0.173±0.004 | -0.192±0.001 | -0.1384±0.0009 |
| **Atenolol** | 2.2[b] | 3.7±0.6 | 3.91±0.03 | 3.7±0.2 | 3.70±0.02 |
| **1-propranolol** | 3.45[c] | 5.6±0.3 | 6.15±0.01 | 5.63±0.09 | 6.31±0.04 |
| **Ibuprofen** | 3.80[c] | 4.9±0.4 | 5.33±0.05 | 4.7±0.2 | 5.65±0.03 |

[a] $LogK_p$ to DMPC, at 25°C, from ref.[4]
[b] $LogK_p$ to soybean PC, at 37°C, from ref.[5]
[c] $LogK_p$ to DOPC, at 25°C, from ref.[6]

**Table S2.** Difference between the logKp for the molecules in the inward and outward directions. Comparison between the values obtained with US and STeUS, after 30 and 100 ns of simulated time.

| Molecule | [$LogK_p$ comp.$^{in}$ , $LogK_p$ comp.$^{out}$] 10 – 30 ns | | 10 – 100 ns | |
|---|---|---|---|---|
| | US | STeUS | US | STeUS |
| **Methanol** | [-0.17, -0.22] | [-0.11, -0.15] | [-0.19, -0.19] | [-0.14, -0.14] |
| **Atenolol** | [1.90, 5.66] | [3.84, 4.00] | [3.03, 4.46] | [3.78, 3.62] |
| **1-propranolol** | [4.56, 6.59] | [6.14, 6.16] | [5.28, 5.98] | [6.48, 6.15] |
| **Ibuprofen** | [3.64, 6.27] | [5.16, 5.50] | [4.12, 5.35] | [5.75, 5.54] |

**Applying the modified STeUS approach yields similar results as standard STeUS**

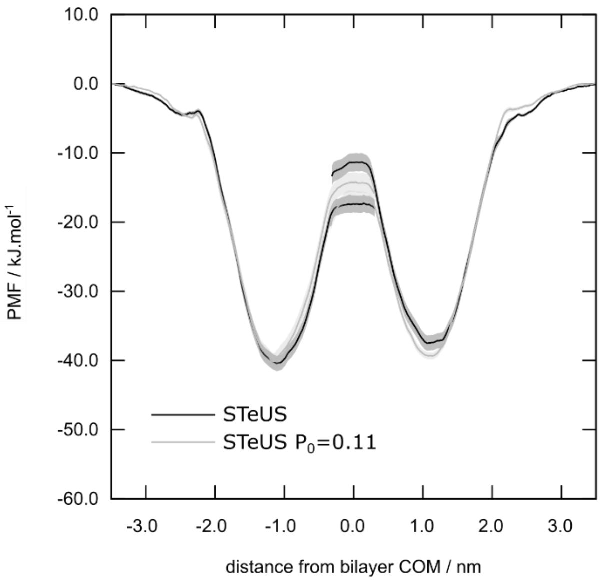


**Figure S5.** PMF of ibuprofen, as a function of solute distance from the bilayer COM, z = 0.0 nm, after 100 ns of simulation time. Solid lines represent the results of two independent simulations for the standard STeUS approach (black line) and the modified STeUS approach with $P_0$= 0.11 (gray). Dashed areas represent the statistical errors, estimated by bootstrapping.

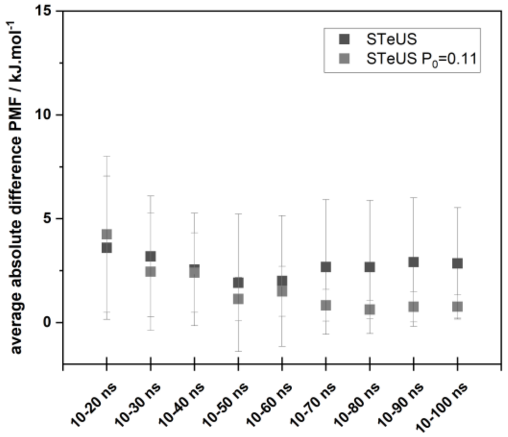


**Figure S6.** Absolute difference between the PMFs in the inward and outward directions of ibuprofen, with increasing simulation times. Data points show the results of two independent simulations for the standard STeUS approach and the modified STeUS approach with $P_0$= 0.11.
