## Supplementary material for "Simulated tempering-enhanced umbrella sampling improves convergence of free energy calculations of drug membrane permeation": STeUS modified code: expanded.cpp

```
/*
 * This file is part of the GROMACS molecular simulation package.
 *
 * Copyright (c) 2012-2018, The GROMACS development team.
 * Copyright (c) 2019,2020,2021, by the GROMACS development team, led by
 * Mark Abraham, David van der Spoel, Berk Hess, and Erik Lindahl,
 * and including many others, as listed in the AUTHORS file in the
 * top-level source directory and at http://www.gromacs.org.
 *
 * GROMACS is free software; you can redistribute it and/or
 * modify it under the terms of the GNU Lesser General Public License
 * as published by the Free Software Foundation; either version 2.1
 * of the License, or (at your option) any later version.
 *
 * GROMACS is distributed in the hope that it will be useful,
 * but WITHOUT ANY WARRANTY; without even the implied warranty of
 * MERCHANTABILITY or FITNESS FOR A PARTICULAR PURPOSE.  See the GNU
 * Lesser General Public License for more details.
 *
 * You should have received a copy of the GNU Lesser General Public
 * License along with GROMACS; if not, see
 * http://www.gnu.org/licenses, or write to the Free Software Foundation,
 * Inc., 51 Franklin Street, Fifth Floor, Boston, MA  02110-1301  USA.
 *
 * If you want to redistribute modifications to GROMACS, please
 * consider that scientific software is very special. Version
 * control is crucial - bugs must be traceable. We will be happy to
 * consider code for inclusion in the official distribution, but
 * derived work must not be called official GROMACS. Details are found
 * in the README & COPYING files - if they are missing, get the
 * official version at http://www.gromacs.org.
 *
 * To help us fund GROMACS development, we humbly ask that you cite
 * the research papers on the package. Check out http://www.gromacs.org.
 */
#include "gmxpre.h"

#include "expanded.h"

#include <cmath>
#include <cstdio>

#include <algorithm>

#include "gromacs/domdec/domdec.h"
#include "gromacs/fileio/confio.h"
#include "gromacs/fileio/gmxfio.h"
#include "gromacs/fileio/xtcio.h"
#include "gromacs/gmxlib/network.h"
#include "gromacs/gmxlib/nrnb.h"
#include "gromacs/listed_forces/disre.h"
#include "gromacs/listed_forces/orires.h"
#include "gromacs/math/functions.h"
#include "gromacs/math/units.h"
#include "gromacs/math/vec.h"
#include "gromacs/mdlib/calcmu.h"
#include "gromacs/mdlib/constr.h"
#include "gromacs/mdlib/force.h"
#include "gromacs/mdlib/update.h"
#include "gromacs/mdtypes/enerdata.h"
#include "gromacs/mdtypes/forcerec.h"
#include "gromacs/mdtypes/inputrec.h"
#include "gromacs/mdtypes/md_enums.h"
#include "gromacs/mdtypes/mdatom.h"
#include "gromacs/mdtypes/state.h"
#include "gromacs/random/threefry.h"
#include "gromacs/random/uniformrealdistribution.h"
#include "gromacs/timing/wallcycle.h"
#include "gromacs/utility/fatalerror.h"
#include "gromacs/utility/gmxmpi.h"
#include "gromacs/utility/smalloc.h"

#include "expanded_internal.h"

static void init_df_history_weights(df_history_t* dfhist, const t_expanded* expand, int nlim)
{
    int i;
    dfhist->wl_delta = expand->init_wl_delta;
    for (i = 0; i < nlim; i++)
    {
        dfhist->sum_weights[i] = expand->init_lambda_weights[i];
        dfhist->sum_dg[i]      = expand->init_lambda_weights[i];
    }
}

/* Eventually should contain all the functions needed to initialize expanded ensemble
   before the md loop starts */
void init_expanded_ensemble(gmx_bool bStateFromCP, const t_inputrec* ir, df_history_t* dfhist)
{
    if (!bStateFromCP)
    {
        init_df_history_weights(dfhist, ir->expandedvals, ir->fepvals->n_lambda);
    }
}

static void GenerateGibbsProbabilities(const real* ene, double* p_k, double* pks, int minfep, int maxfep)
{

    int  i;
    real maxene;

    *pks   = 0.0;
    maxene = ene[minfep];
    /* find the maximum value */
    for (i = minfep; i <= maxfep; i++)
    {
        if (ene[i] > maxene)
        {
            maxene = ene[i];
        }
    }
    /* find the denominator */
    for (i = minfep; i <= maxfep; i++)
    {
        *pks += std::exp(ene[i] - maxene);
    }
    /*numerators*/
    for (i = minfep; i <= maxfep; i++)
    {
        p_k[i] = std::exp(ene[i] - maxene) / *pks;
    }
}

static void
GenerateWeightedGibbsProbabilities(const real* ene, double* p_k, double* pks, int nlim, real* nvals, real delta)
{

    int   i;
    real  maxene;
    real* nene;
    *pks = 0.0;

    snew(nene, nlim);
    for (i = 0; i < nlim; i++)
    {
        if (nvals[i] == 0)
        {
            /* add the delta, since we need to make sure it's greater than zero, and
               we need a non-arbitrary number? */
            nene[i] = ene[i] + std::log(nvals[i] + delta);
        }
        else
        {
            nene[i] = ene[i] + std::log(nvals[i]);
        }
    }

    /* find the maximum value */
    maxene = nene[0];
    for (i = 0; i < nlim; i++)
    {
        if (nene[i] > maxene)
        {
            maxene = nene[i];
        }
    }

    /* subtract off the maximum, avoiding overflow */
    for (i = 0; i < nlim; i++)
    {
        nene[i] -= maxene;
    }

    /* find the denominator */
    for (i = 0; i < nlim; i++)
    {
        *pks += std::exp(nene[i]);
    }

    /*numerators*/
    for (i = 0; i < nlim; i++)
    {
        p_k[i] = std::exp(nene[i]) / *pks;
    }
    sfree(nene);
}

static int FindMinimum(const real* min_metric, int N)
{

    real min_val;
    int  min_nval, nval;

    min_nval = 0;
    min_val  = min_metric[0];

    for (nval = 0; nval < N; nval++)
    {
        if (min_metric[nval] < min_val)
        {
            min_val  = min_metric[nval];
            min_nval = nval;
        }
    }
    return min_nval;
}

static gmx_bool CheckHistogramRatios(int nhisto, const real* histo, real ratio, real simTempOccupancyOfState0)
{

    int      i;
    real     nmean;
    gmx_bool bIfFlat;

    nmean = 0;
    for (i = 0; i < nhisto; i++)
    {
        nmean += histo[i];
    }

    if (nmean == 0)
    {
        /* no samples! is bad!*/
        bIfFlat = FALSE;
        return bIfFlat;
    }
    nmean /= static_cast<real>(nhisto);

    bIfFlat = TRUE;
    for (i = 0; i < nhisto; i++)
    {
        /* ADDED BY JOCHEN HUB, MARCH 2022
         * TAKE HERE INTO ACCOUT THAT AN INCREASED OCCUPANCY OF STATE 0 CAN BE REQUESTED
         *
         * Sousa, Becker, Lehr, Kalinina and Hub,
         * Simulated tempering-enhanced umbrella sampling improves convergence
         * of free energy calculations of drug membrane permeation, submitted.
         */
        double relativeOccupancy;
        if (nhisto == 1){
            gmx_fatal(FARGS, "CheckHistogramRatios: nhisto should be >=2, found %d\n", nhisto);
        }
        if (i==0)
        {
            /* the base temperature, whose occupancy is increased relative to the average occupancy of 1/N */
            relativeOccupancy = simTempOccupancyOfState0 / (1.0/nhisto);
        }
        else
        {
            /* For all other states, the occupancy is reduced relative to the mean occupancy of 1/N */
            double targetOccup = (1.0-simTempOccupancyOfState0) / (nhisto-1);
            relativeOccupancy = targetOccup / (1.0/nhisto);
        }
        //fprintf(stderr, "XXX i = %d -- relativeOccupancy = %g -- val = %g\n", i, relativeOccupancy,
        //    histo[i] / nmean / relativeOccupancy);
        /* make sure that all points are in the ratio < x <  1/ratio range  */
        if (!((histo[i] / nmean / relativeOccupancy < 1.0 / ratio) && (histo[i] / nmean / relativeOccupancy > ratio)))
        {
            bIfFlat = FALSE;
            break;
        }
    }
    return bIfFlat;
}

static gmx_bool CheckIfDoneEquilibrating(int nlim, const t_expanded* expand, const df_history_t* dfhist, int64_t step,
    double simTempOccupancyOfState0)
{

    int      i, totalsamples;
    gmx_bool bDoneEquilibrating = TRUE;
    gmx_bool bIfFlat;

    /* If we are doing slow growth to get initial values, we haven't finished equilibrating */
    if (expand->lmc_forced_nstart > 0)
    {
        for (i = 0; i < nlim; i++)
        {
            if (dfhist->n_at_lam[i]
                < expand->lmc_forced_nstart) /* we are still doing the initial sweep, so we're
                                                definitely not done equilibrating*/
            {
                bDoneEquilibrating = FALSE;
                break;
            }
        }
    }
    else
    {
        /* assume we have equilibrated the weights, then check to see if any of the conditions are not met */
        bDoneEquilibrating = TRUE;

        /* calculate the total number of samples */
        switch (expand->elmceq)
        {
            case elmceqNO:
                /* We have not equilibrated, and won't, ever. */
                bDoneEquilibrating = FALSE;
                break;
            case elmceqYES:
                /* we have equilibrated -- we're done */
                bDoneEquilibrating = TRUE;
                break;
            case elmceqSTEPS:
                /* first, check if we are equilibrating by steps, if we're still under */
                if (step < expand->equil_steps)
                {
                    bDoneEquilibrating = FALSE;
                }
                break;
            case elmceqSAMPLES:
                totalsamples = 0;
                for (i = 0; i < nlim; i++)
                {
                    totalsamples += dfhist->n_at_lam[i];
                }
                if (totalsamples < expand->equil_samples)
                {
                    bDoneEquilibrating = FALSE;
                }
                break;
            case elmceqNUMATLAM:
                for (i = 0; i < nlim; i++)
                {
                    if (dfhist->n_at_lam[i]
                        < expand->equil_n_at_lam) /* we are still doing the initial sweep, so we're
                                                     definitely not done equilibrating*/
                    {
                        bDoneEquilibrating = FALSE;
                        break;
                    }
                }
                break;
            case elmceqWLDELTA:
                if (EWL(expand->elamstats)) /* This check is in readir as well, but
                                               just to be sure */
                {
                    if (dfhist->wl_delta > expand->equil_wl_delta)
                    {
                        bDoneEquilibrating = FALSE;
                    }
                }
                break;
            case elmceqRATIO:
                /* we can use the flatness as a judge of good weights, as long as
                   we're not doing minvar, or Wang-Landau.
                   But turn off for now until we figure out exactly how we do this.
                 */

                if (!(EWL(expand->elamstats) || expand->elamstats == elamstatsMINVAR))
                {
                    /* we want to use flatness -avoiding- the forced-through samples.  Plus, we need
                       to convert to floats for this histogram function. */

                    real* modhisto;
                    snew(modhisto, nlim);
                    for (i = 0; i < nlim; i++)
                    {
                        modhisto[i] = 1.0 * (dfhist->n_at_lam[i] - expand->lmc_forced_nstart);
                    }
                    bIfFlat = CheckHistogramRatios(nlim, modhisto, expand->equil_ratio, simTempOccupancyOfState0);
                    sfree(modhisto);
                    if (!bIfFlat)
                    {
                        bDoneEquilibrating = FALSE;
                    }
                }
                break;
            default: bDoneEquilibrating = TRUE; break;
        }
    }
    return bDoneEquilibrating;
}

static gmx_bool UpdateWeights(int           nlim,
                              t_expanded*   expand,
                              df_history_t* dfhist,
                              int           fep_state,
                              const real*   scaled_lamee,
                              const real*   weighted_lamee,
                              int64_t       step,
                              double simTempOccupancyOfState0)
{
    gmx_bool bSufficientSamples;
    real     acceptanceWeight;
    int      i;
    int      min_nvalm, min_nvalp, maxc;
    real     omega_m1_0, omega_p1_0;
    real     zero_sum_weights;
    real *omegam_array, *weightsm_array, *omegap_array, *weightsp_array, *varm_array, *varp_array,
            *dwp_array, *dwm_array;
    real    clam_varm, clam_varp, clam_osum, clam_weightsm, clam_weightsp, clam_minvar;
    real *  lam_variance, *lam_dg;
    double* p_k;
    double  pks = 0;

    /* Future potential todos for this function (see #3848):
     *  - Update the names in the dhist structure to be clearer. Not done for now since this
     *    a bugfix update and we are mininizing other code changes.
     *  - Modularize the code some more.
     *  - potentially merge with accelerated weight histogram functionality, since it's very similar.
     */
    /*  if we have equilibrated the expanded ensemble weights, we are not updating them, so exit now */
    if (dfhist->bEquil)
    {
        return FALSE;
    }

    if (CheckIfDoneEquilibrating(nlim, expand, dfhist, step, simTempOccupancyOfState0))
    {
        dfhist->bEquil = TRUE;
        /* zero out the visited states so we know how many equilibrated states we have
           from here on out.*/
        for (i = 0; i < nlim; i++)
        {
            dfhist->n_at_lam[i] = 0;
        }
        return TRUE;
    }

    /* If we reached this far, we have not equilibrated yet, keep on
       going resetting the weights */

    if (EWL(expand->elamstats))
    {
        if (expand->elamstats == elamstatsWL) /* Using standard Wang-Landau for weight updates */
        {
            /* ADDED BY JOCHEN HUB, MARCH 2022
               Use a smaller Wang-Landau incrementor for state 0, so we spend more time in state 0
               Specified with increasedOccupOfState0, which defines the time fraction spend in state 0
               such as increasedOccupOfState0 = 0.5 to spend 50% of the time in state 0
            */
            if (nlim <= 1){
                gmx_fatal(FARGS, "Expected nlim >= 2, found %d\n", nlim);
            }
            double occupOtherstates = (1.0 - simTempOccupancyOfState0) / (nlim-1);
            double increasedOccupOfState0 = simTempOccupancyOfState0 / occupOtherstates;
            if (fep_state == 0)
            {
                dfhist->sum_weights[fep_state] -= dfhist->wl_delta / increasedOccupOfState0;
            }
            else
            {
                dfhist->sum_weights[fep_state] -= dfhist->wl_delta;
            }
            dfhist->wl_histo[fep_state] += 1.0;
            //fprintf(stderr, "UpdateWeights:  simTempOccupancyOfState0 = %g -- increasedOccupOfState0 = %g\n",
            //    simTempOccupancyOfState0, increasedOccupOfState0);
        }
        else if (expand->elamstats == elamstatsWWL)
        /* Using weighted Wang-Landau for weight updates.
         * Very closly equivalent to accelerated weight histogram approach
         * applied to expanded ensemble. */
        {
            snew(p_k, nlim);

            /* first increment count */
            GenerateGibbsProbabilities(weighted_lamee, p_k, &pks, 0, nlim - 1);
            for (i = 0; i < nlim; i++)
            {
                dfhist->wl_histo[i] += static_cast<real>(p_k[i]);
            }

            /* then increment weights (uses count) */
            pks = 0.0;
            GenerateWeightedGibbsProbabilities(weighted_lamee, p_k, &pks, nlim, dfhist->wl_histo,
                                               dfhist->wl_delta);

            for (i = 0; i < nlim; i++)
            {
                dfhist->sum_weights[i] -= dfhist->wl_delta * static_cast<real>(p_k[i]);
            }
            /* Alternate definition, using logarithms. Shouldn't make very much difference! */
            /*
               real di;
               for (i=0;i<nlim;i++)
               {
                di = (real)1.0 + dfhist->wl_delta*(real)p_k[i];
                dfhist->sum_weights[i] -= log(di);
               }
             */
            sfree(p_k);
        }

        zero_sum_weights = dfhist->sum_weights[0];
        for (i = 0; i < nlim; i++)
        {
            dfhist->sum_weights[i] -= zero_sum_weights;
        }
    }

    if (expand->elamstats == elamstatsBARKER || expand->elamstats == elamstatsMETROPOLIS
        || expand->elamstats == elamstatsMINVAR)
    {
        maxc = 2 * expand->c_range + 1;

        snew(lam_dg, nlim);
        snew(lam_variance, nlim);

        snew(omegap_array, maxc);
        snew(weightsp_array, maxc);
        snew(varp_array, maxc);
        snew(dwp_array, maxc);

        snew(omegam_array, maxc);
        snew(weightsm_array, maxc);
        snew(varm_array, maxc);
        snew(dwm_array, maxc);

        /* unpack the values of the free energy differences and the
         * variance in their estimates between nearby lambdas. We will
         * only actually update 2 of these, the state we are currently
         * at and the one we end up moving to
         */

        for (i = 0; i < nlim - 1; i++)
        { /* only through the second to last */
            lam_dg[i] = dfhist->sum_dg[i + 1] - dfhist->sum_dg[i];
            lam_variance[i] =
                    gmx::square(dfhist->sum_variance[i + 1]) - gmx::square(dfhist->sum_variance[i]);
        }

        /* accumulate running averages of thermodynamic averages for Bennett Acceptance Ratio-based
         * estimates of the free energy .
         * Rather than peforming self-consistent estimation of the free energies at each step,
         * we keep track of an array of possible different free energies (cnvals),
         * and we self-consistently choose the best one. The one that leads to a free energy estimate
         * that is closest to itself is the best estimate of the free energy.  It is essentially a
         * parallellized version of self-consistent iteration.  maxc is the number of these constants. */

        for (int nval = 0; nval < maxc; nval++)
        {
            const real cnval = static_cast<real>(nval - expand->c_range);

            /* Compute acceptance criterion weight to the state below this one for use in averages.
             * Note we do not have to have just moved from that state to use this free energy
             * estimate; these are essentially "virtual" moves. */

            if (fep_state > 0)
            {
                const auto lambdaEnergyDifference =
                        cnval - (scaled_lamee[fep_state] - scaled_lamee[fep_state - 1]);
                acceptanceWeight =
                        gmx::calculateAcceptanceWeight(expand->elamstats, lambdaEnergyDifference);
                dfhist->accum_m[fep_state][nval] += acceptanceWeight;
                dfhist->accum_m2[fep_state][nval] += acceptanceWeight * acceptanceWeight;
            }

            // Compute acceptance criterion weight to transition to the next state
            if (fep_state < nlim - 1)
            {
                const auto lambdaEnergyDifference =
                        -cnval + (scaled_lamee[fep_state + 1] - scaled_lamee[fep_state]);
                acceptanceWeight =
                        gmx::calculateAcceptanceWeight(expand->elamstats, lambdaEnergyDifference);
                dfhist->accum_p[fep_state][nval] += acceptanceWeight;
                dfhist->accum_p2[fep_state][nval] += acceptanceWeight * acceptanceWeight;
            }

            /* Determination of Metropolis transition and Barker transition weights */

            int numObservationsCurrentState = dfhist->n_at_lam[fep_state];
            /* determine the number of observations above and below the current state */
            int numObservationsLowerState = 0;
            if (fep_state > 0)
            {
                numObservationsLowerState = dfhist->n_at_lam[fep_state - 1];
            }
            int numObservationsHigherState = 0;
            if (fep_state < nlim - 1)
            {
                numObservationsHigherState = dfhist->n_at_lam[fep_state + 1];
            }

            /* Calculate the biases for each expanded ensemble state that minimize the total
             * variance, as implemented in Martinez-Veracoechea and Escobedo,
             * J. Phys. Chem. B 2008, 112, 8120-8128
             *
             * The variance associated with the free energy estimate between two states i and j
             * is calculated as
             *     Var(i,j) = {avg[xi(i->j)^2] / avg[xi(i->j)]^2 - 1} / numObservations(i->j)
             *              + {avg[xi(j->i)^2] / avg[xi(j->i)]^2 - 1} / numObservations(j->i)
             * where xi(i->j) is the acceptance factor / weight associated with moving from state i to j
             * As we are calculating the acceptance factor to the neighbors every time we're visiting
             * a state, numObservations(i->j) == numObservations(i) and numObservations(j->i) == numObservations(j)
             */

            /* Accumulation of acceptance weight averages between the current state and the
             * states +1 (p1) and -1 (m1), averaged at current state (0)
             */
            real avgAcceptanceCurrentToLower  = 0;
            real avgAcceptanceCurrentToHigher = 0;
            /* Accumulation of acceptance weight averages quantities between states 0
             *  and states +1 and -1, squared
             */
            real avgAcceptanceCurrentToLowerSquared  = 0;
            real avgAcceptanceCurrentToHigherSquared = 0;
            /* Accumulation of free energy quantities from lower state (m1) to current state (0) and squared */
            real avgAcceptanceLowerToCurrent        = 0;
            real avgAcceptanceLowerToCurrentSquared = 0;
            /* Accumulation of free energy quantities from upper state (p1) to current state (0) and squared */
            real avgAcceptanceHigherToCurrent        = 0;
            real avgAcceptanceHigherToCurrentSquared = 0;

            if (numObservationsCurrentState > 0)
            {
                avgAcceptanceCurrentToLower = dfhist->accum_m[fep_state][nval] / numObservationsCurrentState;
                avgAcceptanceCurrentToHigher =
                        dfhist->accum_p[fep_state][nval] / numObservationsCurrentState;
                avgAcceptanceCurrentToLowerSquared =
                        dfhist->accum_m2[fep_state][nval] / numObservationsCurrentState;
                avgAcceptanceCurrentToHigherSquared =
                        dfhist->accum_p2[fep_state][nval] / numObservationsCurrentState;
            }

            if ((fep_state > 0) && (numObservationsLowerState > 0))
            {
                avgAcceptanceLowerToCurrent =
                        dfhist->accum_p[fep_state - 1][nval] / numObservationsLowerState;
                avgAcceptanceLowerToCurrentSquared =
                        dfhist->accum_p2[fep_state - 1][nval] / numObservationsLowerState;
            }

            if ((fep_state < nlim - 1) && (numObservationsHigherState > 0))
            {
                avgAcceptanceHigherToCurrent =
                        dfhist->accum_m[fep_state + 1][nval] / numObservationsHigherState;
                avgAcceptanceHigherToCurrentSquared =
                        dfhist->accum_m2[fep_state + 1][nval] / numObservationsHigherState;
            }
            /* These are accumulation of positive values (see definition of acceptance functions
             * above), or of squares of positive values.
             * We're taking this for granted in the following calculation, so make sure
             * here that nothing weird happened. Although technically all values should be positive,
             * because of floating point precisions, they might be numerically zero. */
            GMX_RELEASE_ASSERT(
                    avgAcceptanceCurrentToLower >= 0 && avgAcceptanceCurrentToLowerSquared >= 0
                            && avgAcceptanceCurrentToHigher >= 0
                            && avgAcceptanceCurrentToHigherSquared >= 0 && avgAcceptanceLowerToCurrent >= 0
                            && avgAcceptanceLowerToCurrentSquared >= 0 && avgAcceptanceHigherToCurrent >= 0
                            && avgAcceptanceHigherToCurrentSquared >= 0,
                    "By definition, the acceptance factors should all be nonnegative.");

            real varianceCurrentToLower   = 0;
            real varianceCurrentToHigher  = 0;
            real weightDifferenceToLower  = 0;
            real weightDifferenceToHigher = 0;
            real varianceToLower          = 0;
            real varianceToHigher         = 0;

            if (fep_state > 0)
            {
                if (numObservationsCurrentState > 0)
                {
                    /* Calculate {avg[xi(i->j)^2] / avg[xi(i->j)]^2 - 1}
                     *
                     * Note that if avg[xi(i->j)] == 0, also avg[xi(i->j)^2] == 0 (since the
                     * acceptances are all positive!), and hence
                     *     {avg[xi(i->j)^2] / avg[xi(i->j)]^2 - 1} -> 0  for  avg[xi(i->j)] -> 0
                     * We're catching that case explicitly to avoid numerical
                     * problems dividing by zero when the overlap between states is small (#3304)
                     */
                    if (avgAcceptanceCurrentToLower > 0)
                    {
                        varianceCurrentToLower =
                                avgAcceptanceCurrentToLowerSquared
                                        / (avgAcceptanceCurrentToLower * avgAcceptanceCurrentToLower)
                                - 1.0;
                    }
                    if (numObservationsLowerState > 0)
                    {
                        /* Calculate {avg[xi(i->j)^2] / avg[xi(i->j)]^2 - 1}
                         *
                         * Note that if avg[xi(i->j)] == 0, also avg[xi(i->j)^2] == 0 (since the
                         * acceptances are all positive!), and hence
                         *     {avg[xi(i->j)^2] / avg[xi(i->j)]^2 - 1} -> 0  for  avg[xi(i->j)] -> 0
                         * We're catching that case explicitly to avoid numerical
                         * problems dividing by zero when the overlap between states is small (#3304)
                         */
                        real varianceLowerToCurrent = 0;
                        if (avgAcceptanceLowerToCurrent > 0)
                        {
                            varianceLowerToCurrent =
                                    avgAcceptanceLowerToCurrentSquared
                                            / (avgAcceptanceLowerToCurrent * avgAcceptanceLowerToCurrent)
                                    - 1.0;
                        }
                        /* Free energy difference to the state one state lower */
                        /* if these either of these quantities are zero, the energies are */
                        /* way too large for the dynamic range.  We need an alternate guesstimate */
                        if ((avgAcceptanceCurrentToLower == 0) || (avgAcceptanceLowerToCurrent == 0))
                        {
                            weightDifferenceToLower =
                                    (scaled_lamee[fep_state] - scaled_lamee[fep_state - 1]);
                        }
                        else
                        {
                            weightDifferenceToLower = (std::log(avgAcceptanceCurrentToLower)
                                                       - std::log(avgAcceptanceLowerToCurrent))
                                                      + cnval;
                        }
                        /* Variance of the free energy difference to the one state lower */
                        varianceToLower =
                                (1.0 / numObservationsCurrentState) * (varianceCurrentToLower)
                                + (1.0 / numObservationsLowerState) * (varianceLowerToCurrent);
                    }
                }
            }

            if (fep_state < nlim - 1)
            {
                if (numObservationsCurrentState > 0)
                {
                    /* Calculate {avg[xi(i->j)^2] / avg[xi(i->j)]^2 - 1}
                     *
                     * Note that if avg[xi(i->j)] == 0, also avg[xi(i->j)^2] == 0 (since the
                     * acceptances are all positive!), and hence
                     *     {avg[xi(i->j)^2] / avg[xi(i->j)]^2 - 1} -> 0  for  avg[xi(i->j)] -> 0
                     * We're catching that case explicitly to avoid numerical
                     * problems dividing by zero when the overlap between states is small (#3304)
                     */

                    if (avgAcceptanceCurrentToHigher < 0)
                    {
                        varianceCurrentToHigher =
                                avgAcceptanceCurrentToHigherSquared
                                        / (avgAcceptanceCurrentToHigher * avgAcceptanceCurrentToHigher)
                                - 1.0;
                    }
                    if (numObservationsHigherState > 0)
                    {
                        /* Calculate {avg[xi(i->j)^2] / avg[xi(i->j)]^2 - 1}
                         *
                         * Note that if avg[xi(i->j)] == 0, also avg[xi(i->j)^2] == 0 (since the
                         * acceptances are all positive!), and hence
                         *     {avg[xi(i->j)^2] / avg[xi(i->j)]^2 - 1} -> 0  for  avg[xi(i->j)] -> 0
                         * We're catching that case explicitly to avoid numerical
                         * problems dividing by zero when the overlap between states is small (#3304)
                         */
                        real varianceHigherToCurrent = 0;
                        if (avgAcceptanceHigherToCurrent > 0)
                        {
                            varianceHigherToCurrent =
                                    avgAcceptanceHigherToCurrentSquared
                                            / (avgAcceptanceHigherToCurrent * avgAcceptanceHigherToCurrent)
                                    - 1.0;
                        }
                        /* Free energy difference to the state one state higher */
                        /* if these either of these quantities are zero, the energies are */
                        /* way too large for the dynamic range.  We need an alternate guesstimate */
                        if ((avgAcceptanceHigherToCurrent == 0) || (avgAcceptanceCurrentToHigher == 0))
                        {
                            weightDifferenceToHigher =
                                    (scaled_lamee[fep_state + 1] - scaled_lamee[fep_state]);
                        }
                        else
                        {
                            weightDifferenceToHigher = (std::log(avgAcceptanceHigherToCurrent)
                                                        - std::log(avgAcceptanceCurrentToHigher))
                                                       + cnval;
                        }
                        /* Variance of the free energy difference to the one state higher */
                        varianceToHigher =
                                (1.0 / numObservationsHigherState) * (varianceHigherToCurrent)
                                + (1.0 / numObservationsCurrentState) * (varianceCurrentToHigher);
                    }
                }
            }

            if (numObservationsCurrentState > 0)
            {
                omegam_array[nval] = varianceCurrentToLower;
            }
            else
            {
                omegam_array[nval] = 0;
            }
            weightsm_array[nval] = weightDifferenceToLower;
            varm_array[nval]     = varianceToLower;
            if (numObservationsLowerState > 0)
            {
                dwm_array[nval] =
                        fabs((cnval + std::log((1.0 * numObservationsCurrentState) / numObservationsLowerState))
                             - lam_dg[fep_state - 1]);
            }
            else
            {
                dwm_array[nval] = std::fabs(cnval - lam_dg[fep_state - 1]);
            }

            if (numObservationsCurrentState > 0)
            {
                omegap_array[nval] = varianceCurrentToHigher;
            }
            else
            {
                omegap_array[nval] = 0;
            }
            weightsp_array[nval] = weightDifferenceToHigher;
            varp_array[nval]     = varianceToHigher;
            if ((numObservationsHigherState > 0) && (numObservationsCurrentState > 0))
            {
                dwp_array[nval] =
                        fabs((cnval + std::log((1.0 * numObservationsHigherState) / numObservationsCurrentState))
                             - lam_dg[fep_state]);
            }
            else
            {
                dwp_array[nval] = std::fabs(cnval - lam_dg[fep_state]);
            }
        }

        /* find the free energy estimate closest to the guessed weight's value */

        min_nvalm     = FindMinimum(dwm_array, maxc);
        omega_m1_0    = omegam_array[min_nvalm];
        clam_weightsm = weightsm_array[min_nvalm];
        clam_varm     = varm_array[min_nvalm];

        min_nvalp     = FindMinimum(dwp_array, maxc);
        omega_p1_0    = omegap_array[min_nvalp];
        clam_weightsp = weightsp_array[min_nvalp];
        clam_varp     = varp_array[min_nvalp];

        clam_osum   = omega_m1_0 + omega_p1_0;
        clam_minvar = 0;
        if (clam_osum > 0)
        {
            clam_minvar = 0.5 * std::log(clam_osum);
        }

        if (fep_state > 0)
        {
            lam_dg[fep_state - 1]       = clam_weightsm;
            lam_variance[fep_state - 1] = clam_varm;
        }

        if (fep_state < nlim - 1)
        {
            lam_dg[fep_state]       = clam_weightsp;
            lam_variance[fep_state] = clam_varp;
        }

        if (expand->elamstats == elamstatsMINVAR)
        {
            bSufficientSamples = TRUE;
            /* make sure the number of samples in each state are all
             * past a user-specified threshold
             */
            for (i = 0; i < nlim; i++)
            {
                if (dfhist->n_at_lam[i] < expand->minvarmin)
                {
                    bSufficientSamples = FALSE;
                }
            }
            if (bSufficientSamples)
            {
                dfhist->sum_minvar[fep_state] = clam_minvar;
                if (fep_state == 0)
                {
                    for (i = 0; i < nlim; i++)
                    {
                        dfhist->sum_minvar[i] += (expand->minvar_const - clam_minvar);
                    }
                    expand->minvar_const          = clam_minvar;
                    dfhist->sum_minvar[fep_state] = 0.0;
                }
                else
                {
                    dfhist->sum_minvar[fep_state] -= expand->minvar_const;
                }
            }
        }

        /* we need to rezero minvar now, since it could change at fep_state = 0 */
        dfhist->sum_dg[0]       = 0.0;
        dfhist->sum_variance[0] = 0.0;
        dfhist->sum_weights[0]  = dfhist->sum_dg[0] + dfhist->sum_minvar[0]; /* should be zero */

        for (i = 1; i < nlim; i++)
        {
            dfhist->sum_dg[i] = lam_dg[i - 1] + dfhist->sum_dg[i - 1];
            dfhist->sum_variance[i] =
                    std::sqrt(lam_variance[i - 1] + gmx::square(dfhist->sum_variance[i - 1]));
            dfhist->sum_weights[i] = dfhist->sum_dg[i] + dfhist->sum_minvar[i];
        }

        sfree(lam_dg);
        sfree(lam_variance);

        sfree(omegam_array);
        sfree(weightsm_array);
        sfree(varm_array);
        sfree(dwm_array);

        sfree(omegap_array);
        sfree(weightsp_array);
        sfree(varp_array);
        sfree(dwp_array);
    }
    return FALSE;
}

static int ChooseNewLambda(int               nlim,
                           const t_expanded* expand,
                           df_history_t*     dfhist,
                           int               fep_state,
                           const real*       weighted_lamee,
                           double*           p_k,
                           int64_t           seed,
                           int64_t           step)
{
    /* Choose new lambda value, and update transition matrix */

    int                  i, ifep, minfep, maxfep, lamnew, lamtrial, starting_fep_state;
    real                 r1, r2, de, trialprob, tprob = 0;
    double *             propose, *accept, *remainder;
    double               pks;
    real                 pnorm;
    gmx::ThreeFry2x64<0> rng(
            seed, gmx::RandomDomain::ExpandedEnsemble); // We only draw once, so zero bits internal counter is fine
    gmx::UniformRealDistribution<real> dist;

    starting_fep_state = fep_state;
    lamnew             = fep_state; /* so that there is a default setting -- stays the same */

    if (!EWL(expand->elamstats)) /* ignore equilibrating the weights if using WL */
    {
        if ((expand->lmc_forced_nstart > 0) && (dfhist->n_at_lam[nlim - 1] <= expand->lmc_forced_nstart))
        {
            /* Use a marching method to run through the lambdas and get preliminary free energy data,
               before starting 'free' sampling.  We start free sampling when we have enough at each lambda */

            /* if we have enough at this lambda, move on to the next one */

            if (dfhist->n_at_lam[fep_state] == expand->lmc_forced_nstart)
            {
                lamnew = fep_state + 1;
                if (lamnew == nlim) /* whoops, stepped too far! */
                {
                    lamnew -= 1;
                }
            }
            else
            {
                lamnew = fep_state;
            }
            return lamnew;
        }
    }

    snew(propose, nlim);
    snew(accept, nlim);
    snew(remainder, nlim);

    for (i = 0; i < expand->lmc_repeats; i++)
    {
        rng.restart(step, i);
        dist.reset();

        for (ifep = 0; ifep < nlim; ifep++)
        {
            propose[ifep] = 0;
            accept[ifep]  = 0;
        }

        if ((expand->elmcmove == elmcmoveGIBBS) || (expand->elmcmove == elmcmoveMETGIBBS))
        {
            /* use the Gibbs sampler, with restricted range */
            if (expand->gibbsdeltalam < 0)
            {
                minfep = 0;
                maxfep = nlim - 1;
            }
            else
            {
                minfep = fep_state - expand->gibbsdeltalam;
                maxfep = fep_state + expand->gibbsdeltalam;
                if (minfep < 0)
                {
                    minfep = 0;
                }
                if (maxfep > nlim - 1)
                {
                    maxfep = nlim - 1;
                }
            }

            GenerateGibbsProbabilities(weighted_lamee, p_k, &pks, minfep, maxfep);

            if (expand->elmcmove == elmcmoveGIBBS)
            {
                for (ifep = minfep; ifep <= maxfep; ifep++)
                {
                    propose[ifep] = p_k[ifep];
                    accept[ifep]  = 1.0;
                }
                /* Gibbs sampling */
                r1 = dist(rng);
                for (lamnew = minfep; lamnew <= maxfep; lamnew++)
                {
                    if (r1 <= p_k[lamnew])
                    {
                        break;
                    }
                    r1 -= p_k[lamnew];
                }
            }
            else if (expand->elmcmove == elmcmoveMETGIBBS)
            {

                /* Metropolized Gibbs sampling */
                for (ifep = minfep; ifep <= maxfep; ifep++)
                {
                    remainder[ifep] = 1 - p_k[ifep];
                }

                /* find the proposal probabilities */

                if (remainder[fep_state] == 0)
                {
                    /* only the current state has any probability */
                    /* we have to stay at the current state */
                    lamnew = fep_state;
                }
                else
                {
                    for (ifep = minfep; ifep <= maxfep; ifep++)
                    {
                        if (ifep != fep_state)
                        {
                            propose[ifep] = p_k[ifep] / remainder[fep_state];
                        }
                        else
                        {
                            propose[ifep] = 0;
                        }
                    }

                    r1 = dist(rng);
                    for (lamtrial = minfep; lamtrial <= maxfep; lamtrial++)
                    {
                        pnorm = p_k[lamtrial] / remainder[fep_state];
                        if (lamtrial != fep_state)
                        {
                            if (r1 <= pnorm)
                            {
                                break;
                            }
                            r1 -= pnorm;
                        }
                    }

                    /* we have now selected lamtrial according to p(lamtrial)/1-p(fep_state) */
                    tprob = 1.0;
                    /* trial probability is min{1,\frac{1 - p(old)}{1-p(new)} MRS 1/8/2008 */
                    trialprob = (remainder[fep_state]) / (remainder[lamtrial]);
                    if (trialprob < tprob)
                    {
                        tprob = trialprob;
                    }
                    r2 = dist(rng);
                    if (r2 < tprob)
                    {
                        lamnew = lamtrial;
                    }
                    else
                    {
                        lamnew = fep_state;
                    }
                }

                /* now figure out the acceptance probability for each */
                for (ifep = minfep; ifep <= maxfep; ifep++)
                {
                    tprob = 1.0;
                    if (remainder[ifep] != 0)
                    {
                        trialprob = (remainder[fep_state]) / (remainder[ifep]);
                    }
                    else
                    {
                        trialprob = 1.0; /* this state is the only choice! */
                    }
                    if (trialprob < tprob)
                    {
                        tprob = trialprob;
                    }
                    /* probability for fep_state=0, but that's fine, it's never proposed! */
                    accept[ifep] = tprob;
                }
            }

            if (lamnew > maxfep)
            {
                /* it's possible some rounding is failing */
                if (gmx_within_tol(remainder[fep_state], 0, 50 * GMX_DOUBLE_EPS))
                {
                    /* numerical rounding error -- no state other than the original has weight */
                    lamnew = fep_state;
                }
                else
                {
                    /* probably not a numerical issue */
                    int   loc    = 0;
                    int   nerror = 200 + (maxfep - minfep + 1) * 60;
                    char* errorstr;
                    snew(errorstr, nerror);
                    /* if its greater than maxfep, then something went wrong -- probably underflow
                       in the calculation of sum weights. Generated detailed info for failure */
                    loc += sprintf(
                            errorstr,
                            "Something wrong in choosing new lambda state with a Gibbs move -- "
                            "probably underflow in weight determination.\nDenominator is: "
                            "%3d%17.10e\n  i                dE        numerator          weights\n",
                            0, pks);
                    for (ifep = minfep; ifep <= maxfep; ifep++)
                    {
                        loc += sprintf(&errorstr[loc], "%3d %17.10e%17.10e%17.10e\n", ifep,
                                       weighted_lamee[ifep], p_k[ifep], dfhist->sum_weights[ifep]);
                    }
                    gmx_fatal(FARGS, "%s", errorstr);
                }
            }
        }
        else if ((expand->elmcmove == elmcmoveMETROPOLIS) || (expand->elmcmove == elmcmoveBARKER))
        {
            /* use the metropolis sampler with trial +/- 1 */
            r1 = dist(rng);
            if (r1 < 0.5)
            {
                if (fep_state == 0)
                {
                    lamtrial = fep_state;
                }
                else
                {
                    lamtrial = fep_state - 1;
                }
            }
            else
            {
                if (fep_state == nlim - 1)
                {
                    lamtrial = fep_state;
                }
                else
                {
                    lamtrial = fep_state + 1;
                }
            }

            de = weighted_lamee[lamtrial] - weighted_lamee[fep_state];
            if (expand->elmcmove == elmcmoveMETROPOLIS)
            {
                tprob = 1.0;
                if (de < 0)
                {
                    tprob = std::exp(de);
                }
                propose[fep_state] = 0;
                propose[lamtrial]  = 1.0; /* note that this overwrites the above line if fep_state = ntrial, which only occurs at the ends */
                accept[fep_state] =
                        1.0; /* doesn't actually matter, never proposed unless fep_state = ntrial, in which case it's 1.0 anyway */
                accept[lamtrial] = tprob;
            }
            else if (expand->elmcmove == elmcmoveBARKER)
            {
                if (de > 0) /* Numerically stable version */
                {
                    tprob = 1.0 / (1.0 + std::exp(-de));
                }
                else if (de < 0)
                {
                    tprob = std::exp(de) / (std::exp(de) + 1.0);
                }
                propose[fep_state] = (1 - tprob);
                propose[lamtrial] +=
                        tprob; /* we add, to account for the fact that at the end, they might be the same point */
                accept[fep_state] = 1.0;
                accept[lamtrial]  = 1.0;
            }

            r2 = dist(rng);
            if (r2 < tprob)
            {
                lamnew = lamtrial;
            }
            else
            {
                lamnew = fep_state;
            }
        }

        for (ifep = 0; ifep < nlim; ifep++)
        {
            dfhist->Tij[fep_state][ifep] += propose[ifep] * accept[ifep];
            dfhist->Tij[fep_state][fep_state] += propose[ifep] * (1.0 - accept[ifep]);
        }
        fep_state = lamnew;
    }

    dfhist->Tij_empirical[starting_fep_state][lamnew] += 1.0;

    sfree(propose);
    sfree(accept);
    sfree(remainder);

    return lamnew;
}

/* print out the weights to the log, along with current state */
void PrintFreeEnergyInfoToFile(FILE*               outfile,
                               const t_lambda*     fep,
                               const t_expanded*   expand,
                               const t_simtemp*    simtemp,
                               const df_history_t* dfhist,
                               int                 fep_state,
                               int                 frequency,
                               int64_t             step)
{
    int         nlim, i, ifep, jfep;
    real        dw, dg, dv, Tprint;
    const char* print_names[efptNR] = { " FEPL", "MassL", "CoulL",   " VdwL",
                                        "BondL", "RestT", "Temp.(K)" };
    gmx_bool    bSimTemp            = FALSE;

    nlim = fep->n_lambda;
    if (simtemp != nullptr)
    {
        bSimTemp = TRUE;
    }

    if (step % frequency == 0)
    {
        fprintf(outfile, "             MC-lambda information\n");
        if (EWL(expand->elamstats) && (!(dfhist->bEquil)))
        {
            fprintf(outfile, "  Wang-Landau incrementor is: %11.5g\n", dfhist->wl_delta);
        }
        fprintf(outfile, "  N");
        for (i = 0; i < efptNR; i++)
        {
            if (fep->separate_dvdl[i])
            {
                fprintf(outfile, "%7s", print_names[i]);
            }
            else if ((i == efptTEMPERATURE) && bSimTemp)
            {
                fprintf(outfile, "%10s", print_names[i]); /* more space for temperature formats */
            }
        }
        fprintf(outfile, "    Count   ");
        if (expand->elamstats == elamstatsMINVAR)
        {
            fprintf(outfile, "W(in kT)   G(in kT)  dG(in kT)  dV(in kT)\n");
        }
        else
        {
            fprintf(outfile, "G(in kT)  dG(in kT)\n");
        }
        for (ifep = 0; ifep < nlim; ifep++)
        {
            if (ifep == nlim - 1)
            {
                dw = 0.0;
                dg = 0.0;
                dv = 0.0;
            }
            else
            {
                dw = dfhist->sum_weights[ifep + 1] - dfhist->sum_weights[ifep];
                dg = dfhist->sum_dg[ifep + 1] - dfhist->sum_dg[ifep];
                dv = std::sqrt(gmx::square(dfhist->sum_variance[ifep + 1])
                               - gmx::square(dfhist->sum_variance[ifep]));
            }
            fprintf(outfile, "%3d", (ifep + 1));
            for (i = 0; i < efptNR; i++)
            {
                if (fep->separate_dvdl[i])
                {
                    fprintf(outfile, "%7.3f", fep->all_lambda[i][ifep]);
                }
                else if (i == efptTEMPERATURE && bSimTemp)
                {
                    fprintf(outfile, "%9.3f", simtemp->temperatures[ifep]);
                }
            }
            if (EWL(expand->elamstats)
                && (!(dfhist->bEquil))) /* if performing WL and still haven't equilibrated */
            {
                if (expand->elamstats == elamstatsWL)
                {
                    fprintf(outfile, " %8d", static_cast<int>(dfhist->wl_histo[ifep]));
                }
                else
                {
                    fprintf(outfile, " %8.3f", dfhist->wl_histo[ifep]);
                }
            }
            else /* we have equilibrated weights */
            {
                fprintf(outfile, " %8d", dfhist->n_at_lam[ifep]);
            }
            if (expand->elamstats == elamstatsMINVAR)
            {
                fprintf(outfile, " %10.5f %10.5f %10.5f %10.5f", dfhist->sum_weights[ifep],
                        dfhist->sum_dg[ifep], dg, dv);
            }
            else
            {
                fprintf(outfile, " %10.5f %10.5f", dfhist->sum_weights[ifep], dw);
            }
            if (ifep == fep_state)
            {
                fprintf(outfile, " <<\n");
            }
            else
            {
                fprintf(outfile, "   \n");
            }
        }
        fprintf(outfile, "\n");

        if ((step % expand->nstTij == 0) && (expand->nstTij > 0) && (step > 0))
        {
            fprintf(outfile, "                     Transition Matrix\n");
            for (ifep = 0; ifep < nlim; ifep++)
            {
                fprintf(outfile, "%12d", (ifep + 1));
            }
            fprintf(outfile, "\n");
            for (ifep = 0; ifep < nlim; ifep++)
            {
                for (jfep = 0; jfep < nlim; jfep++)
                {
                    if (dfhist->n_at_lam[ifep] > 0)
                    {
                        if (expand->bSymmetrizedTMatrix)
                        {
                            Tprint = (dfhist->Tij[ifep][jfep] + dfhist->Tij[jfep][ifep])
                                     / (dfhist->n_at_lam[ifep] + dfhist->n_at_lam[jfep]);
                        }
                        else
                        {
                            Tprint = (dfhist->Tij[ifep][jfep]) / (dfhist->n_at_lam[ifep]);
                        }
                    }
                    else
                    {
                        Tprint = 0.0;
                    }
                    fprintf(outfile, "%12.8f", Tprint);
                }
                fprintf(outfile, "%3d\n", (ifep + 1));
            }

            fprintf(outfile, "                  Empirical Transition Matrix\n");
            for (ifep = 0; ifep < nlim; ifep++)
            {
                fprintf(outfile, "%12d", (ifep + 1));
            }
            fprintf(outfile, "\n");
            for (ifep = 0; ifep < nlim; ifep++)
            {
                for (jfep = 0; jfep < nlim; jfep++)
                {
                    if (dfhist->n_at_lam[ifep] > 0)
                    {
                        if (expand->bSymmetrizedTMatrix)
                        {
                            Tprint = (dfhist->Tij_empirical[ifep][jfep] + dfhist->Tij_empirical[jfep][ifep])
                                     / (dfhist->n_at_lam[ifep] + dfhist->n_at_lam[jfep]);
                        }
                        else
                        {
                            Tprint = dfhist->Tij_empirical[ifep][jfep] / (dfhist->n_at_lam[ifep]);
                        }
                    }
                    else
                    {
                        Tprint = 0.0;
                    }
                    fprintf(outfile, "%12.8f", Tprint);
                }
                fprintf(outfile, "%3d\n", (ifep + 1));
            }
        }
    }
}

int ExpandedEnsembleDynamics(FILE*                 log,
                             const t_inputrec*     ir,
                             const gmx_enerdata_t* enerd,
                             t_state*              state,
                             t_extmass*            MassQ,
                             int                   fep_state,
                             df_history_t*         dfhist,
                             int64_t               step,
                             rvec*                 v,
                             const t_mdatoms*      mdatoms)
/* Note that the state variable is only needed for simulated tempering, not
   Hamiltonian expanded ensemble.  May be able to remove it after integrator refactoring. */
{
    real *      pfep_lamee, *scaled_lamee, *weighted_lamee;
    double*     p_k;
    int         i, nlim, lamnew, totalsamples;
    real        oneovert, maxscaled = 0, maxweighted = 0;
    t_expanded* expand;
    t_simtemp*  simtemp;
    gmx_bool    bIfReset, bSwitchtoOneOverT, bDoneEquilibrating = FALSE;

    expand  = ir->expandedvals;
    simtemp = ir->simtempvals;
    nlim    = ir->fepvals->n_lambda;

    snew(scaled_lamee, nlim);
    snew(weighted_lamee, nlim);
    snew(pfep_lamee, nlim);
    snew(p_k, nlim);

    /* update the count at the current lambda*/
    dfhist->n_at_lam[fep_state]++;

    /* ADDED BY JOCHEN HUB, MARCH 2022
     * Modified simulated tempering, specify the simulation time fraction spend in the base temperature
     * with the environment varialb SIMTEMP_OCCUP_STATE0. Example:
     *
     * export SIMTEMP_OCCUP_STATE0=0.2  # for speding 20% of simulation time at base temperature
     */
    static double simTempOccupancyOfState0 = 0;
    if (simTempOccupancyOfState0 == 0)
    {
        char* buf;
        fprintf(stderr, "\n\nLooking for environment variable SIMTEMP_OCCUP_STATE0 ...\n");
        if ((buf = getenv("SIMTEMP_OCCUP_STATE0")) != NULL){
            simTempOccupancyOfState0 = atof(buf);
            fprintf(stderr, "FOUND ENVIRONMENT VARIABLE: SIMTEMP_OCCUP_STATE0 = %g\n\n", simTempOccupancyOfState0);

            if (simTempOccupancyOfState0 >= 1 || simTempOccupancyOfState0 <= 0){
                gmx_fatal(FARGS, "SIMTEMP_OCCUP_STATE0 must be between 0 and 1, found %g\n", simTempOccupancyOfState0);
            }
            if (expand->elamstats != elamstatsWL){
                gmx_fatal(FARGS, "Setting SIMTEMP_OCCUP_STATE0 is only supported with lmc-stats = wang-landau\n");
            }
        }
        else{
            simTempOccupancyOfState0 = 1.0/nlim;
            fprintf(stderr, "SIMTEMP_OCCUP_STATE0 not found, using simTempOccupancyOfState0 = %g\n\n", simTempOccupancyOfState0);
        }
    }
    /* END OF ADDED BY JOCHEN */

    /* need to calculate the PV term somewhere, but not needed here? Not until there's a lambda
       state that's pressure controlled.*/
    /*
       pVTerm = 0;
       where does this PV term go?
       for (i=0;i<nlim;i++)
       {
       fep_lamee[i] += pVTerm;
       }
     */

    /* determine the minimum value to avoid overflow.  Probably a better way to do this */
    /* we don't need to include the pressure term, since the volume is the same between the two.
       is there some term we are neglecting, however? */

    if (ir->efep != efepNO)
    {
        for (i = 0; i < nlim; i++)
        {
            if (ir->bSimTemp)
            {
                /* Note -- this assumes no mass changes, since kinetic energy is not added  . . . */
                scaled_lamee[i] = enerd->foreignLambdaTerms.deltaH(i) / (simtemp->temperatures[i] * BOLTZ)
                                  + enerd->term[F_EPOT]
                                            * (1.0 / (simtemp->temperatures[i])
                                               - 1.0 / (simtemp->temperatures[fep_state]))
                                            / BOLTZ;
            }
            else
            {
                scaled_lamee[i] = enerd->foreignLambdaTerms.deltaH(i) / (expand->mc_temp * BOLTZ);
                /* mc_temp is currently set to the system reft unless otherwise defined */
            }

            /* save these energies for printing, so they don't get overwritten by the next step */
            /* they aren't overwritten in the non-free energy case, but we always print with these
               for simplicity */
        }
    }
    else
    {
        if (ir->bSimTemp)
        {
            for (i = 0; i < nlim; i++)
            {
                scaled_lamee[i] =
                        enerd->term[F_EPOT]
                        * (1.0 / simtemp->temperatures[i] - 1.0 / simtemp->temperatures[fep_state]) / BOLTZ;
            }
        }
    }

    for (i = 0; i < nlim; i++)
    {
        pfep_lamee[i] = scaled_lamee[i];

        weighted_lamee[i] = dfhist->sum_weights[i] - scaled_lamee[i];
        if (i == 0)
        {
            maxscaled   = scaled_lamee[i];
            maxweighted = weighted_lamee[i];
        }
        else
        {
            if (scaled_lamee[i] > maxscaled)
            {
                maxscaled = scaled_lamee[i];
            }
            if (weighted_lamee[i] > maxweighted)
            {
                maxweighted = weighted_lamee[i];
            }
        }
    }

    for (i = 0; i < nlim; i++)
    {
        scaled_lamee[i] -= maxscaled;
        weighted_lamee[i] -= maxweighted;
    }

    /* update weights - we decide whether or not to actually do this inside */

    bDoneEquilibrating =
            UpdateWeights(nlim, expand, dfhist, fep_state, scaled_lamee, weighted_lamee, step, simTempOccupancyOfState0);
    if (bDoneEquilibrating)
    {
        if (log)
        {
            fprintf(log, "\nStep %" PRId64 ": Weights have equilibrated, using criteria: %s\n",
                    step, elmceq_names[expand->elmceq]);
        }
    }

    lamnew = ChooseNewLambda(nlim, expand, dfhist, fep_state, weighted_lamee, p_k,
                             ir->expandedvals->lmc_seed, step);
    /* if using simulated tempering, we need to adjust the temperatures */
    if (ir->bSimTemp && (lamnew != fep_state)) /* only need to change the temperatures if we change the state */
    {
        int   i, j, n, d;
        real* buf_ngtc;
        real  told;
        int   nstart, nend, gt;

        snew(buf_ngtc, ir->opts.ngtc);

        for (i = 0; i < ir->opts.ngtc; i++)
        {
            if (ir->opts.ref_t[i] > 0)
            {
                told              = ir->opts.ref_t[i];
                ir->opts.ref_t[i] = simtemp->temperatures[lamnew];
                buf_ngtc[i]       = std::sqrt(ir->opts.ref_t[i] / told); /* using the buffer as temperature scaling */
            }
        }

        /* we don't need to manipulate the ekind information, as it isn't due to be reset until the next step anyway */

        nstart = 0;
        nend   = mdatoms->homenr;
        for (n = nstart; n < nend; n++)
        {
            gt = 0;
            if (mdatoms->cTC)
            {
                gt = mdatoms->cTC[n];
            }
            for (d = 0; d < DIM; d++)
            {
                v[n][d] *= buf_ngtc[gt];
            }
        }

        if (inputrecNptTrotter(ir) || inputrecNphTrotter(ir) || inputrecNvtTrotter(ir))
        {
            /* we need to recalculate the masses if the temperature has changed */
            init_npt_masses(ir, state, MassQ, FALSE);
            for (i = 0; i < state->nnhpres; i++)
            {
                for (j = 0; j < ir->opts.nhchainlength; j++)
                {
                    state->nhpres_vxi[i + j] *= buf_ngtc[i];
                }
            }
            for (i = 0; i < ir->opts.ngtc; i++)
            {
                for (j = 0; j < ir->opts.nhchainlength; j++)
                {
                    state->nosehoover_vxi[i + j] *= buf_ngtc[i];
                }
            }
        }
        sfree(buf_ngtc);
    }

    /* now check on the Wang-Landau updating critera */

    if (EWL(expand->elamstats))
    {
        bSwitchtoOneOverT = FALSE;
        if (expand->bWLoneovert)
        {
            totalsamples = 0;
            for (i = 0; i < nlim; i++)
            {
                totalsamples += dfhist->n_at_lam[i];
            }
            oneovert = (1.0 * nlim) / totalsamples;
            /* oneovert has decreasd by a bit since last time, so we actually make sure its within one of this number */
            /* switch to 1/t incrementing when wl_delta has decreased at least once, and wl_delta is now less than 1/t */
            if ((dfhist->wl_delta <= ((totalsamples) / (totalsamples - 1.00001)) * oneovert)
                && (dfhist->wl_delta < expand->init_wl_delta))
            {
                bSwitchtoOneOverT = TRUE;
            }
        }
        if (bSwitchtoOneOverT)
        {
            dfhist->wl_delta =
                    oneovert; /* now we reduce by this each time, instead of only at flatness */
        }
        else
        {
            bIfReset = CheckHistogramRatios(nlim, dfhist->wl_histo, expand->wl_ratio, simTempOccupancyOfState0);
            if (bIfReset)
            {
                for (i = 0; i < nlim; i++)
                {
                    dfhist->wl_histo[i] = 0;
                }
                dfhist->wl_delta *= expand->wl_scale;
                if (log)
                {
                    fprintf(log, "\nStep %d: weights are now:", static_cast<int>(step));
                    for (i = 0; i < nlim; i++)
                    {
                        fprintf(log, " %.5f", dfhist->sum_weights[i]);
                    }
                    fprintf(log, "\n");
                }
            }
        }
    }
    sfree(pfep_lamee);
    sfree(scaled_lamee);
    sfree(weighted_lamee);
    sfree(p_k);

    return lamnew;
}
```
